# Recurrent selection explains parallel evolution of genomic regions of high relative but low absolute differentiation in greenish warblers

**DOI:** 10.1101/041467

**Authors:** Darren E. Irwin, Miguel Alcaide, Kira E. Delmore, Jessica H. Irwin, Gregory L. Owens

## Abstract

Recent technological developments allow investigation of the repeatability of evolution at the genomic level. Such investigation is particularly powerful when applied to a ring species, in which spatial variation represents changes during the evolution of two species from one. We examined genomic variation among three subspecies of the greenish warbler ring species, using genotypes at 13,013,950 nucleotide sites along a new greenish warbler consensus genome assembly. Genomic regions of low within-group variation are remarkably consistent between the three populations. These regions show high relative differentiation but low absolute differentiation between populations. Comparisons with outgroup species show the locations of these peaks of relative differentiation are not well explained by phylogenetically-conserved variation in recombination rates or selection. These patterns are consistent with a model in which selection in an ancestral form has reduced variation at some parts of the genome, and those same regions experience recurrent selection that subsequently reduces variation within each subspecies. The degree of heterogeneity in nucleotide diversity is greater than explained by models background selection, but are consistent with selective sweeps. Given the evidence that greenish warblers have had both population differentiation for a long period of time and periods of gene flow between those populations, we propose that some genomic regions underwent selective sweeps over a broad geographic area followed by within-population selection-induced reductions in variation. An important implication of this “sweep-before-differentiation” model is that genomic regions of high relative differentiation may have moved among populations more recently than other genomic regions.

## Introduction

The question of “How repeatable is evolution?” has captured the interest and imagination of generations of scientists, and motivated much empirical and theoretical research (e.g., Gould 1990; Travisano et al. 1995; Wichman et al. 1999; Wood et al. 2005; Conte et al. 2012; Meyer et al. 2012; Renaut et al. 2014; Bauer and Gokhale 2015). Thought experiments and empirical investigations of this question have mostly focused on phenotypic patterns and/or genetic changes in particular genes of interest, and answers appear to depend on the timescale considered. Considering long timescales in the history of life, Gould (1990) wrote that “any replay of the tape would lead evolution down a pathway radically different from the road actually taken.” On a much shorter timeframe, Lenski and others (e.g., Meyer et al. 2010) have shown strikingly parallel evolution in replicate laboratory populations of bacteria. On intermediate timescales, groups such as stickleback fish (Colosimo et al. 2005) and sunflowers (Renaut et al. 2014) show strong patterns of parallel evolutionary responses to similar environments.

The rapid development of genomic technology now allows expansion of investigations of the repeatability of evolution to DNA sequences across the whole-genome scale (Lobkovsky and Koonin 2012). When an ancestral species evolves into several differentiated descendent populations, do similar regions of the genome appear to play a key role in differentiation? Alternatively, are the patterns highly unrelated, with little similarity between daughter populations in the regions that display differentiation? Investigations of such questions are most powerful when they involve more than just two populations. Here, we investigate patterns of genomic differentiation in a ring species (Mayr 1942; Cain 1954; Irwin et al. 2001c), a situation that allows comparison of the structuring of genomic differentiation at a range of spatial and temporal scales. In a ring species, two terminal forms are reproductively isolated (to a large degree) where they co-occur, but these forms are connected by a long chain of populations encircling an uninhabited area; through this chain there is a gradient in phenotypic and genetic traits and little if any reproductive isolation. We examine patterns of genomic differentiation between the two terminal forms as well as between each of them and a population halfway along the chain connecting them, and we ask how similar the patterns of differentiation are between the three comparisons.

The ring species under investigation is the greenish warbler (*Phylloscopus trochiloides*) species complex (Fig. 1), which consists of two forms breeding in Siberia (*P. t. viridanus* in the west, and *P. t. plumbeitarsus* in the east) and a connecting chain of three subspecies to the south that form a gradient around the uninhabited Tibetan Plateau (*P. t. ludlowi* in the western Himalayas, *P. t. trochiloides* in the central and eastern Himalayas, and *P. t. obscuratus* in central China; Fig. 1; Ticehurst 1938; Mayr 1942). Previous genetic and phenotypic analysis (Irwin 2000, 2012; Irwin et al. 2001b, 2005, 2008; Alcaide et al. 2014) has indicated that there is strong (but not complete) reproductive isolation between *viridanus* and *plumbeitarsus* where they meet in central Siberia, whereas around the southern ring there is little reproductive isolation, although there are indications of some phases of geographic separation followed by secondary contact. The geographic history of the complex is likely very complicated, given the Pleistocene history of many phases of glaciation cycles, but it is clear from the genetic and phenotypic data that west Siberian *viridanus* expanded into Siberia from central Asia (i.e. from the western side of the current ring) and east Siberian *plumbeitarsus* expanded into Siberia from eastern Asia (i.e. from the east side of the current ring).

**Fig. 1.**
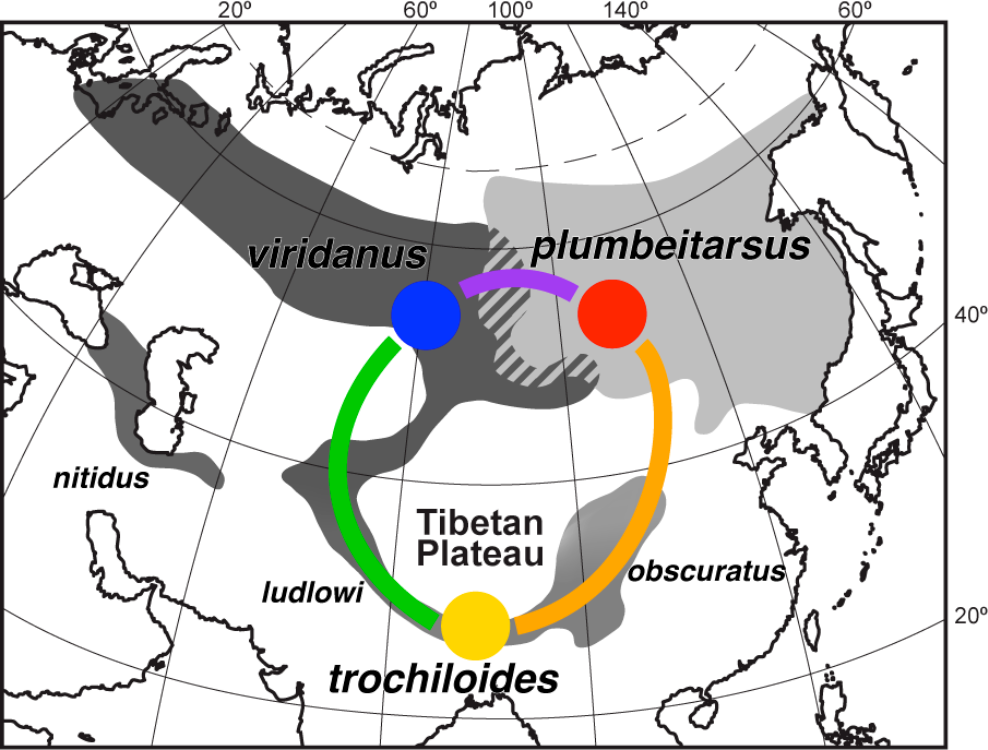
Map of the greenish warbler range, indicating the three main populations under study, and the comparisons between them. The gradient around the greenish warbler ring is shown in shades of gray, and colors indicate the populations (circles) and population comparisons (lines) shown in subsequent figures (note the green and orange line colors represent population comparisons, not other populations as in previous publications). Names of subspecies are indicated, with the three being compared in larger font.

Phenotypic variation around the ring indicates that there has been a combination of parallel and divergent evolution in different traits during the two northward expansions. Body size, seasonal migration distance, and song length have evolved in parallel, with the two Siberian forms (*viridanus* and *plumbeitarsus*) showing strong similarity to each other and both differing in the same way from the southern forms; parallel evolution in these traits is likely due to parallel shifts in habitat and other environmental characteristics during the two northward expansions. In contrast, plumage patterning, migratory routes, and song and call structure have evolved strong differences between the Siberian forms. Some of these differences (e.g. plumage and structure of vocalizations) are likely due to the complexities of sexual and social selection in causing highly stochastic patterns of evolution (Irwin 2000, 2012; Irwin et al. 2001b, 2008). Hence, based on phenotypic patterns, we have reason to expect some parallel and some divergent selection on the genome.

A number of studies of genomic differentiation between pairs of populations have observed distinct chromosomal regions with much higher relative differentiation (i.e., *F*_ST_) than most of the genome (e.g., flycatchers, Ellegren et al. 2012, Burri et al. 2015; mosquitos, Turner et al. 2005; rabbits, Carneiro et al. 2014; mice, Harr 2006; butterflies, Nadeau et al. 2012); these regions have been referred to as “genomic islands of speciation,” “genomic islands of differentiation,” and “genomic islands of divergence.” Two primary explanations have been given for such regions (Nachman and Payseur 2012; Cruickshank and Hahn 2014; Fig. 2A,B). First, in a context of speciation with gene flow (Fig. 2A), the islands of high relative differentiation form because they contain loci involved in reproductive isolation (i.e., “speciation genes”) within the hybrid zone between the two populations, causing those loci to have low gene flow between the two populations compared to other parts of the genome (Wu 2001; Nosil et al. 2009; Feder and Nosil 2010; Nosil and Feder 2012; Via 2012). Loci in high physical linkage with those speciation genes undergo hitchhiking with the speciation genes, such that an island of differentiation forms, facilitating the buildup of linked loci that contribute further to reproductive isolation. Second, in a model of selection in allopatry (Fig. 2B), islands of high relative differentiation are not directly caused by loci causing reproductive isolation when gene flow is occurring, but rather by loci under background and/or directional selection in one or both populations (Noor and Bennett 2009; Turner and Hahn 2010; Nachman and Payseur 2012; Cruickshank and Hahn 2014; Delmore et al. 2015). The selection causes reduced within-population variation at the selected locus as well as areas in close physical linkage; reduced within-population variation tends to be accompanied by greater relative differentiation between populations, since relative differentiation is generally estimated as a ratio of between-group nucleotide differentiation to the total nucleotide variation (the sum of between-group and within-group nucleotide variation).

**Fig. 2.**
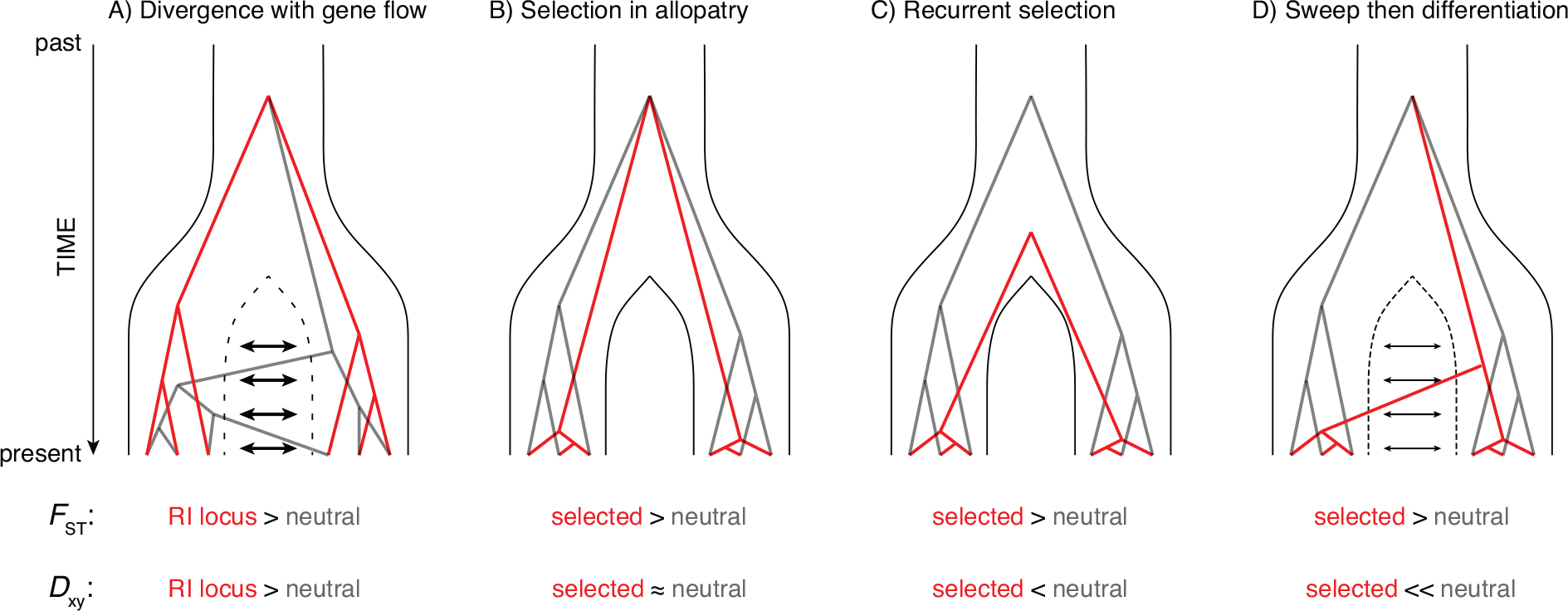
Depictions of four explanations of how genomic areas of high relative differentiation develop, along with comparisons of the expected levels of relative (*F*_ST_) and absolute (*D*_xy_) differentiation between selected and neutral parts of the genome. Each panel depicts a history in which a single ancestral population split into two daughter taxa. In A and D there is some amount of migration between the two populations, whereas in B and C there is not. In each scenario, typical genealogies of six individuals (three in each taxon) are shown for a selected region (in red; in A, “RI locus” refers to a locus causing reproductive isolation) and a neutral locus (in grey).

In both models, close physical linkage of genes plays an important role, because linkage reduces recombination between selected loci and nearby neutral loci, preserving the association between a particular set of alleles (Feder and Nosil 2010; Nachman and Payseur 2012). Hence, there is expected to be a relationship between islands of differentiation and areas of low recombination, as observed in sunflowers (Renaut et al. 2013) and flycatchers (Burri et al. 2015). Just as recombination rates vary across the genome, intensity of selection is expected to vary across the genome, and the combined effect of recombination and selection together influence the amount of heterogeneity in differentiation across the genome (Nachman and Payseur 2012; Renaut et al. 2013; Burri et al. 2015).

Nachman and Payseur (2012)and Cruickshank and Hahn (2014)proposed a way to distinguish the standard speciation-with-gene-flow model and the selection-in-allopatry model for the formation of peaks of high relative differentiation, by examining patterns of absolute nucleotide differentiation (i.e., *D*_xy_) between groups, rather than focusing primarily on patterns of relative differentiation (i.e., *F*_ST_). Under the standard speciation-with-gene-flow model (Fig. 2A), absolute nucleotide differentiation is expected to be high in the islands of high relative differentiation, because reproductive isolation due to loci in those regions prevents those regions from flowing between populations, whereas the rest of the genome flows between populations, eventually reducing absolute differentiation. In contrast, under a selection-in-allopatry model (Fig. 2B), absolute differentiation is not expected to be high where relative differentiation is high; rather, increased relative differentiation in areas with loci under selection is due solely to decreased within-group variation (i.e., π). Hence, in our comparisons of patterns of genomic differentiation around the greenish warbler ring, we compare patterns of within-and between-group nucleotide variation as well as relative differentiation.

Our focus here is on describing and understanding the causes of patterns of genomic heterogeneity in nucleotide diversity and and differentiation among three populations of greenish warblers, rather than on questions of overall relationships of populations around the ring, which was the focus of previous work (e.g., Alcaide et al. 2014). We ask several specific questions. First, are there distinct genomic regions of high relative differentiation between populations? Second, how similar are regions of differentiation in the three pairwise population comparisons? Given that there are two distinct south-to-north geographic clines in greenish warblers (west and east of the Tibetan Plateau), we can compare the patterns of genomic differentiation that have occurred along each. Third, do regions of high relative differentiation display high absolute differentiation (supporting the speciation-with-gene-flow model for the formation of such high-*F*_ST_regions)? Alternatively, is high relative differentiation due entirely to low within-group variation (supporting the selection-in-allopatry model)? To test whether patterns of nucleotide variation across the genome may be explained by variation in mutation rate or recombination rate, we also analyze genomic variation in four outgroup taxa. Finally, we ask whether the Z chromosome (a sex chromosome) shows different levels of within- and between-population variation compared to autosomes, since theory and previous observational studies in other systems suggest that sex chromosomes may differentiate faster and play an especially important role in speciation (Charlesworth et al. 1987; Ellegren et al. 2012; reviewed by Oyler-McCance et al. 2015).

## Material and Methods

### Sampling

We used DNA extracts obtained from blood samples of wild-caught birds. These included 135 samples broadly distributed around the greenish warbler ring; broad genomic relationships among these were previously summarized in Alcaide et al. (2014). We also included in the present study four samples of other species related to greenish warblers (*Phylloscopus inornatus, Phylloscopus humei, Seicercus whistleri*, and *Phylloscopus fuscatus*). Of these, *S. whistleri*is most closely related to greenish warblers (despite having a different genus name), and the other three are in a different clade, with *P. inornatus*and *P. humei*being sister taxa (Johansson et al. 2007; Price 2010).

### Building a Greenish Warbler Consensus Reference Genome

For the purpose of mapping GBS reads to a reference genome, we wished to construct a reference genome based on variation throughout the greenish warbler ring, such that our mapping of genetic variation would not be biased toward any one part of the ring. Hence we chose one individual from each of the three most divergent subspecies around the ring (*viridanus*, bird TL2; *trochiloides*, LN10; and *plumbeitarsus*, BK2), conducted whole-genome shotgun sequencing and de novo assembly on each, and then constructed a consensus reference sequence based on those three individuals. Each of the three whole-genome sequencing libraries (one for each reference individual) were run within one lane of an Illumina HiSeq 2000 automated sequencer at the NextGen Sequencing Facility of the Biodiversity Research Centre (University of British Columbia, Vancouver, Canada).

We trimmed and removed duplicates from each library using Trimmomatic (version 0.32; Bolger et al. 2014), using the settings “TRAILING:3 SLIDINGWINDOW:4:10 MINLEN:30” and FastUniq (version 1.1, default settings, Xu et al. 2012), respectively. The Avian Genome Consortium (Zhang et al. 2014) recently assembled genomes for 48 birds using SOAPdenovo (version 1.05, kmer of 27, Luo et al. 2012). We used the same settings to obtain de novo assemblies for each of our three greenish warblers (k=27, d=1 and M=3; https://github.com/gigascience/paper-zhang2014/tree/master/Genome_assembly/S0APdenovo). Summary statistics for the resulting assemblies are presented in Table S1. The length of each assembly is similar to other avian genomes (between 1 and 1.2 Gb; Ellegren 2013). Faircloth et al. (2012)identified two sets of ultra-conserved elements (UCEs) using whole genome alignments for the chicken, anole and zebra finch. The first set included 5561 elements; the second was limited to UCEs with higher coverage and included 2560 elements. We aligned these sequences to each genome using NCBI’s blastn. Results are shown in Table S1 and show that all assemblies had at least 91 % of the first set and 99.5% of the second set.

To order the scaffolds in each de novo assembly and organize into putative chromosomes, we used BWA to align scaffolds to the repeat-masked version of the zebra finch genome (version 3.2.4), resulting in a high fraction of scaffolds mapping (TL2: 78.4%; LN10: 82.4%; BK2: 80.2%); given the high synteny of the avian genome between chicken and zebra finch (Warren et al. 2010; Ellegren 2013; Kawakami et al. 2014), we make the assumption that synteny is also high between zebra finch and the warblers studied here. We imported these alignments into Geneious, which was used to construct the reference sequence for each chromosome as follows. For each individual, we annotated all regions of the zebra finch reference that had no coverage in the greenish warbler sequences, and then we extracted the consensus sequence, using a consensus threshold of 0% (fewest ambiguities), and with “If no coverage call Ref” checked. This resulted in a reference genome for each individual, consisting of greenish warbler sequence where there was coverage, and zebra finch sequence where there was not coverage (but with those no-coverage regions annotated). We used the Mauve plugin (with the ProgressiveMauve algorithm on default settings, except with “Assume collinear genomes” checked; Darling et al. 2010) to align the three individual references to each other. This Mauve alignment was then extracted, and then all regions within each reference sequence that had no greenish warbler sequence coverage (that is, those annotated as a no-coverage region) were converted to missing bases. We then trimmed the ends of the alignments consisting of only missing bases, and then extracted the consensus of these three sequences (using consensus threshold of 0%, with “ignore gaps” checked, and “If no coverage call N”). This procedure resulted in a reference sequence for each chromosome consisting only of consensus greenish warbler sequence, but of similar length as the zebra finch reference chromosome.

For the great majority of chromosomal regions, the alignment steps in the above procedure appeared to work very well, but for a few small regions (Table S2) the initial alignments in the Mauve step appeared poor and too long, with many large gaps inserted in each sequence such that very little sequence in each individual actually aligned to sequence in other individuals. For these regions, an additional alignment step was added, involving realigning that small region with customized parameters (e.g. adjusting the “seed weight” in Mauve). In every case, a parameter set was found that resulted in a good (and much shorter) alignment.

### Mapping of Gbs Reads and Genotyping

Raw GBS reads from Alcaide et al. (2014)were re-analyzed using an entirely distinct bioinformatic pipeline. These reads were produced by paired-end Illumina sequencing of two libraries, the first containing 96 samples broadly distributed around the greenish warbler ring, and the second containing 70 samples, 65 of which (39 adults, 25 chicks, and 1 duplicate sample for control purposes) were from a single research site (Keylong; site code PA) along the southwest side of the ring (the other 5 were outgroup species, only 4 of which produced good sequence; see “Groups in each analysis” below). Alcaide et al. (2014)describes details of library preparation and numbers of GBS reads produced (briefly, roughly 3.3 million reads per individual in the first library, and 4.5 million in the second library).

Reads were de-multiplexed according to the in-line GBS barcode using a custom Perl script that separated reads, removed barcode and adaptor sequence, and removed sequences shorter than 30 bp in length. The de-multiplexing script allowed no mismatches in barcode sequence., Reads were then trimmed for base quality using Trimmomatic-0.32 (with options TRAILING:3 SLIDINGWINDOW:4:10 MINLEN:30). We used BWA-MEM (Li and Durbin 2009) on default settings to align trimmed reads to our greenish warbler consensus genome, and the programs Picard (http://broadinstitute.github.io/picard/) and SAMtools (Li et al. 2009) were used to produce BAM files containing the alignments. The program GATK (McKenna *et al.*2010) was then used to realign reads around indels (using the tools *RealignerTargetCreator*, followed by *IndelRealigner*) and then call genotypes (*HaplotypeCaller*, with options “--emitRefConfidence GVCF--max_alternate_alleles 2-variant_index_type LINEAR-variant_index_parameter 128000”), resulting in a GVCF file for each individual. Genotyping information from all individuals within an analysis was then combined into a single file for each chromosome using the GATK command *GenotypeGVCFs*, with the option “-allSites” used such that genotypes at both variant and invariant sites were retained, and the option “-L” used to specify the chromosome (data was separated by chromosome at this point, in order to reduce file size and facilitate downstream computational efficiency).

A variety of analyses limited to different sets of individuals were then performed (see below), with the GATK command *SelectVariants*used to choose information for each set of individuals. We then used a combination of VCFtools (Danecek et al. 2011) and custom-written scripts to apply a series of filters to determine which sites were included in the analysis: First, indels and SNPs with more than 2 alleles were removed, to avoid the complicating effects of such variants on the calculation of differentiation statistics. Second, we removed sites with more than 40% of individuals had missing genotypes, to restrict the analysis to sites with data from a substantial portion of individuals. Third, we removed sites with MQ < 20, to avoid poorly mapped reads. Fourth, we removed sites with heterozygosity above 60%, to avoid paralogs. We converted the resulting vcf file to a matrix of genotypes of each individual at each site.

### Illustration of Genomic Relationships Around the Ring

We used principal coordinates analysis (using custom scripts in R version 3.1.2, employing the “pca” command with method “svdImpute” to account for missing genotypes) to summarize and visualize genomic relationships among individuals. We first filtered out any individuals that were missing genotypes at more than 25% of the SNPs identified in the set of individuals included in a particular analysis. We centered but did not scale genotypic values, thereby ensuring that each nucleotide mismatch had equal weighting in the analysis (this means that more variable SNPs have larger influence in the analysis).

### Estimation of Differentiation Statistics Across the Genome

A custom script in R (R Core Team 2014) was used to estimate summary differentiation statistics and to produce graphs of variation across the genome. First, for each nucleotide site we calculated allele frequencies for each group of individuals defined in an analysis. We used these frequencies to calculate, for each site, both within-and between-group average pairwise differentiation between two individuals. Within-group nucleotide differentiation was calculated as 2p(1−p), where p is the frequency of one of the alleles (each nucleotide had either 1 [invariant] or 2 alleles); this within-group nucleotide differentiation thus ranges from zero to 0.5. Between-group nucleotide differentiation was calculated as p_1_(1−p_2_) + p_2_(1−p_1_), where p_1_is the frequency of a given allele in the first group and p_2_is the frequency of that allele in the second group; this ranges from zero to one.

For sites that were variable, we estimated nucleotide-specific *F*_ST_according to Weir and Cockerham’s (1984) equation for 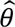(top of their page 1363). This method assumes random mating within populations, and corrects for two types of sampling bias due to limited sample size: that due to limited sample size of individuals within groups, and that due to sampling a limited number of populations out of all possible replicate populations (both real and imagined, under the same evolutionary parameters that the real sampled populations evolved). We used this method to estimate *F*_ST_for each nucleotide for each pair of populations included in an analysis, and also to estimate *F*_ST_among all groups in an analysis. For sample sizes, we used the numbers of individuals successfully genotyped at that specific nucleotide site in that specific population.

Given our focus on patterns of differentiation across the genome rather than at individual nucleotide sites, we also calculated averages of these statistics on windows across each chromosome. In order to ensure that summary statistics for each window were not influenced by sample size of nucleotide sites within each window, windows were defined based on a fixed number of sites (5000 or 10,000 nucleotides depending on the analysis; see next section) for which we had good genotypic information (that is, they survived the filtering process described above), rather than a fixed window size across the reference genome. Windows were aligned starting from the side of each chromosome corresponding to the beginning side of that chromosome in the zebra finch genome (i.e., the left side in figures), and summary statistics were not calculated for incomplete window fragments on the other (right) side.

For each window, we calculated mean within-group nucleotide differentiation (π) for each group, and mean between-group nucleotide differentiation (*D*_xy_) for each pair of groups; these statistics incorporate information from both variant and invariant sites. We also estimated multilocus *F*_ST_ for each window by summing the numerators of the 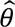equation across sites and then dividing by the sum of the denominators of the same equation across sites (Weir and Cockerham 1984).

### Groups in Each Analysis

Our analysis focused on two major analyses of differentiation within and between populations. In the first, we included 15 individuals from each of the three major greenish warbler taxa (*viridanus, trochiloides, plumbeitarsus*; Table S3a). Individuals were chosen for inclusion in these groups prior to examining the genotypic data, and were based on choosing individuals with both high sequence coverage and that well represent the core population of that taxon (i.e., far from known hybrid zones, and avoiding those individuals identified by Alcaide et al. [2014] as having some chromosome fragments from other taxa). We view this sampling procedure as appropriate because we wanted our analysis to represent differentiation between the core populations of each taxon.

In the second analysis, we included one individual from each of nine taxa (five greenish warblers, of which three are analyzed here; and four outgroup species; Table S4). While this is certainly a small sample of each taxon, note (i) that each individual is diploid and thus contains one of each chromosome from each parent, meaning the sample size of chromosomes of each taxon is two, and (ii) a windowed analysis summarizes patterns at thousands of nucleotides in each window, reducing the impact of sampling error on windowed averages. However, because of the increased sampling error compared to the 15-individual-per-taxon analysis above, and because heterozygosity of individuals tends to be underestimated using low-coverage sequencing data, the magnitude of differentiation statistics should not be compared directly between the two analyses. Nevertheless, overall patterns of variation within and among greenish warbler populations were very similar across chromosomes in the two analyses, indicating that our comparisons with outgroup species are also valid, and data from one individual per taxon was sufficient to recover strong correlations in within-group variation observed across the genome between *viridanus, trochiloides*, and *viridanus*using larger sample sizes (Fig. S2).

Window size was set at 5000 nucleotide sites in the first (45-sample) analysis (corresponding to an average of 144 SNPs per window), and 10,000 in the second (9-sample) analysis (due to the greater influence of noise in the second).

## Results

Mapping of GBS reads to the reference greenish warbler genome resulted in the identification of 580,356 single nucleotide polymorphisms among 135 greenish warbler samples. Whole-genome relationships, as summarized using principal components analysis (PCA; Fig. S1), show the pattern expected based on previous research (Alcaide et al. 2014;see also Bradburd et al. 2016), of two highly distinct Siberian forms (*viridanus*and *plumbeitarsus*) and a progression of genomic signatures through the ring of populations to the south. Note that Alcaide et al. (2014)summarized variation in the same GBS reads, but used an entirely distinct bioinformatics pipeline and based their PCA on only 2,334 SNPs due to very restrictive filtering; the fact that the current study recovers similar patterns using much less restrictive filtering and roughly 250 times the number of SNPs gives strong confidence in the inferred relationships. We use these overall genomic relationships as a backdrop to explore patterns of variation in relative and absolute differentiation across the genome.

We first examined differentiation between three major geographic groups around the ring: *viridanus* in west Siberia, *trochiloides* in the south, and *plumbeitarsus* in east Siberia. To ensure that variation in sample size did not influence comparisons of patterns among these taxa, we chose 15 individuals of each taxon to include in an analysis of between-group relative nucleotide differentiation (*F*_ST_), between-group absolute nucleotide differentiation (*D*_xy_), and within-group nucleotide variation (π). To estimate absolute nucleotide differentiation, we used our GBS reads to identify invariant nucleotide sites as well as variant ones, resulting in a dataset of 12,639,111 invariant and 374,839 variant nucleotide sites among our 45 samples in the analysis, and calculated statistics in windows across each chromosome. We show results first for a single chromosome (Fig. 3), and then for the entire genome (Fig. 4).

**Fig. 3.**
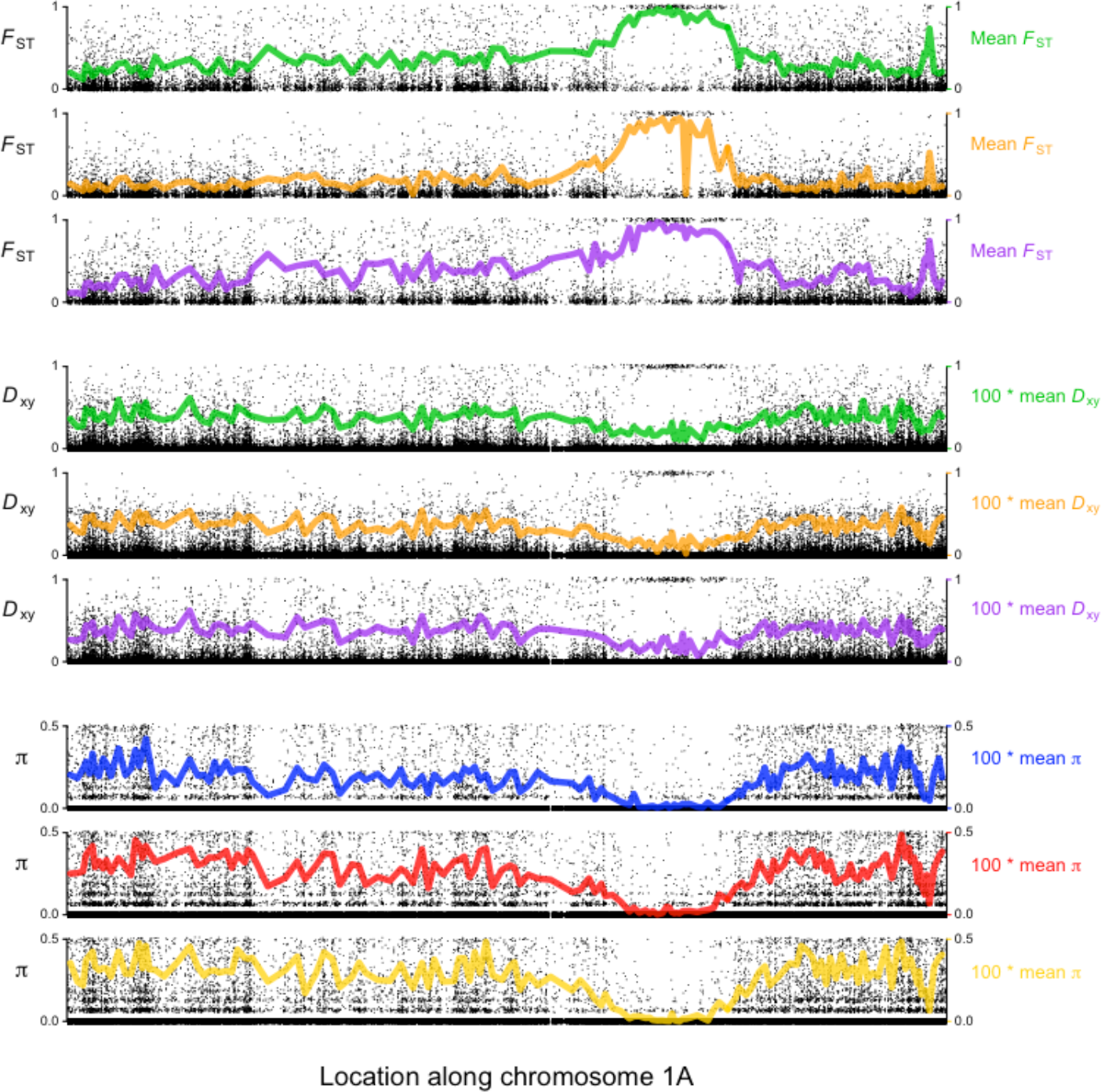
Nucleotide differentiation across chromosome 1A shows a region of consistently strong relative differentiation (*F*_ST_, top) and low absolute differentiation (*D*_xy_, middle) between greenish warbler populations, as well as extremely low within-population variation (π, bottom). Each graph shows per-nucleotide statistics for 647,363 nucleotides, 24,764 of which are variable among a dataset of 15 *viridanus*, 15 *trochiloides*, and 15 *plumbeitarsus*, and colored lines show windowed averages (5000 nucleotide sites per window; averages for *D*_xy_and π are multiplied by 100, to better use the vertical axis). The top three graphs show *F*_ST_(only defined for variable markers) between *trochiloides*and *viridanus*(top, green), *trochiloides*and *plumbeitarsus*(middle, orange), and *viridanus*and *plumbeitarsus*(bottom, purple). The middle three graphs show *D*_xy_, using the same population comparisons as above. The lower three graphs show π within *viridanus*(top, blue), *plumbeitarsus*(middle, red), and trochiloides (bottom, yellow). Note that a small amount of jittering (2.5% of each vertical axis) was added to individual nucleotide statistics (i.e., the black dots).

**Fig. 4.**
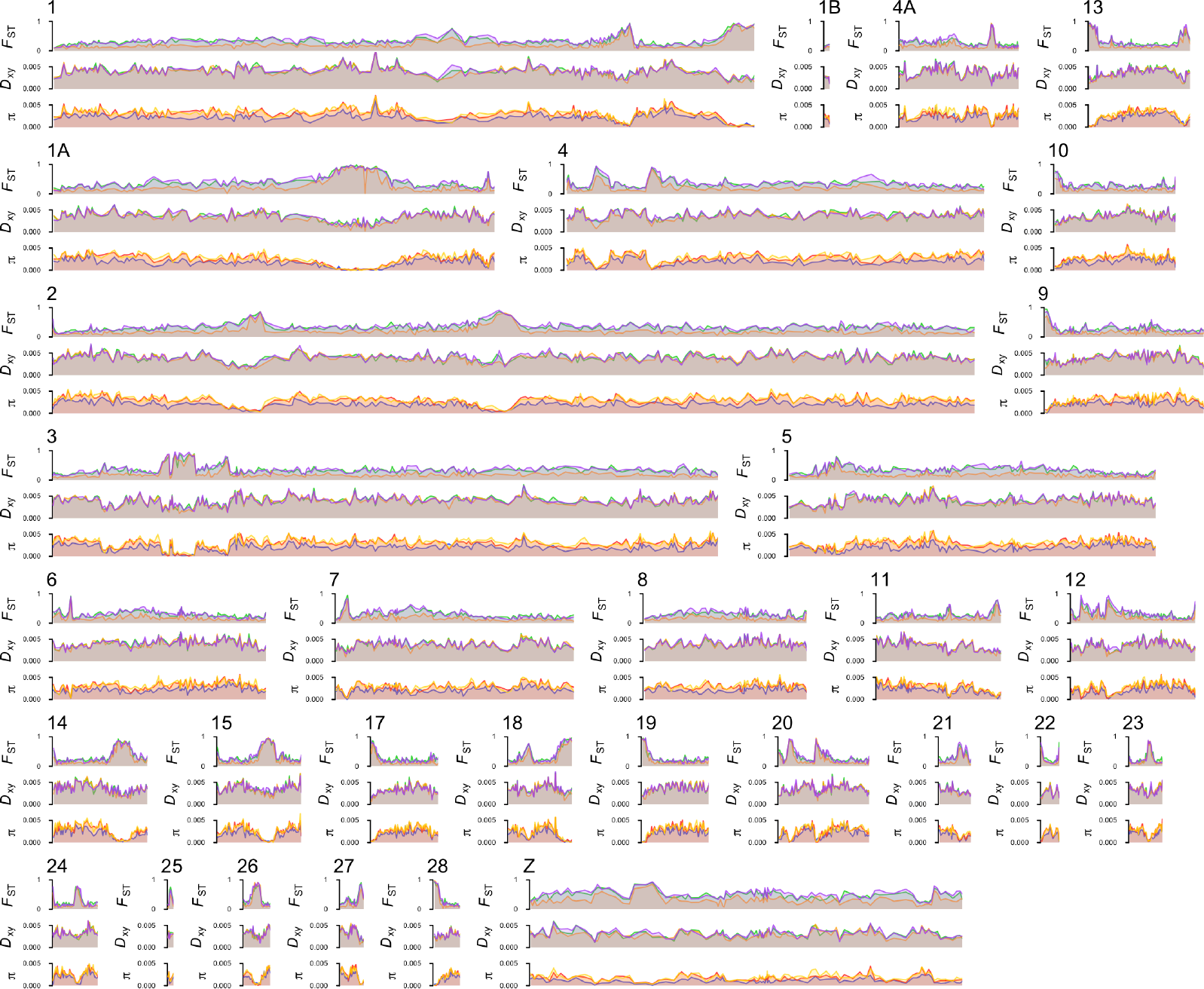
Nucleotide differentiation shows consistent patterns of strong genomic structuring within and between greenish warbler populations. For each chromosome, graphs show variation in per-window relative nucleotide differentiation (*F*_ST_, top), absolute nucleotide differentiation (*D*_xy_, middle), and within-group nucleotide variation (π, bottom), using the same colors (3 often overlapping lines for each small plot) for particular population comparisons (for *F*_ST_ and *D*_xy_) and populations (for π) as in Figs. 1 and 3.

Relative differentiation shows tremendous variability across the genome, with most chromosomes having one to several distinct islands of high relative differentiation against a background of much lower relative differentiation. Remarkably, the locations of peaks of high relative differentiation are highly similar in all pairwise comparisons among the three taxa (Figs. 3–5; see statistical tests in caption of Fig. 5). The Z chromosome on average shows much higher levels of relative differentiation than the autosomes (e.g., mean *F*_ST_ among windows in the comparison of *viridanus* and *plumbeitarsus*: 0.33 for autosomes vs. 0.52 for Z chromosome; Welch’s t-test: t = −12.1, df = 115.9, P < 10^−15^); for this reason we focus on autosomes below, and return to the Z-chromosome later.

**Fig. 5.**
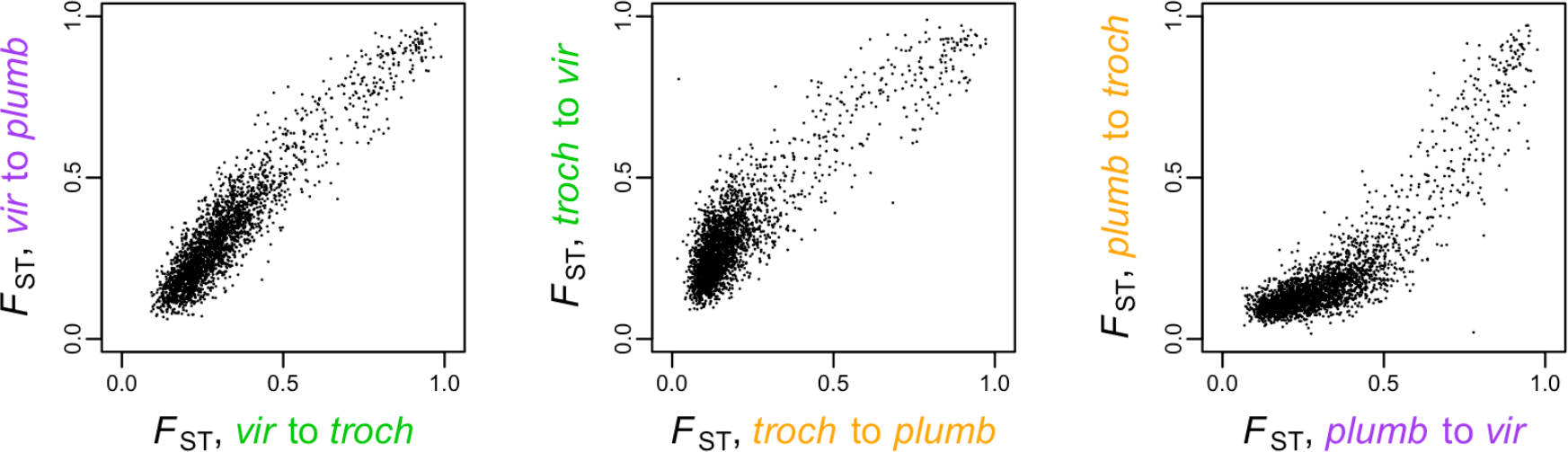
Per-window relative differentiation (*F*_ST_) between each pair of greenish warbler populations is strongly correlated with that between each of the other pairs. Each graph is a bivariate plot of *F*_ST_ between one pair of populations vs. *F*_ST_ between a second pair, with each dot representing one of 2486 windows across the autosomal genome, each consisting of 5000 nucleotide sites. Within each comparison, there is a dense low-*F*_ST_ cluster containing the great majority of windows, and a long string of higher-*F*_ST_ windows. These tend to have high *F*_ST_ in all three comparisons (Spearman’s rank correlations: *vir* to *troch* vs. *vir* to *plumb*: *r_s_* = 0.876, P < 10 ^−15^; *troch* to *plumb* vs *troch* to *vir*: *r_s_* = 0.697, P < 10^−15^; *plumb* to *vir* vs. *plumb* to *troch*: *r*_s_ = 0.760, P < 10^−15^).

Surprisingly, regions of high relative differentiation usually have low absolute differentiation (Figs. 3,4,6), and this is consistent among all comparisons (Spearman’s rank correlations: *trochiloides*vs. *viridanus*: P < 9.4*10^−9^; *trochiloides* vs. *plumbeitarsus*: P < 10^−15^; *viridanus* vs. *plumbeitarsus*: P = 0.0096; *n* = 2486 windows). What accounts for this apparent paradox? It is the remarkably low within-group nucleotide diversity (π) in these genomic regions (Fig. 7). These regions of low within-group diversity have strikingly similar locations in all three taxa (Figs. 3,4,8a). Moreover, these regions have much lower within-group diversity than would be proportional to the reduced between-group absolute differentiation alone: in Fig. 8b we show that regions with a low ratio of within-group diversity to between-group absolute differentiation, which we call “standardized nucleotide diversity,” are consistent among all three greenish warbler taxa. Hence, the regions of high relative differentiation have lower within-and between-group absolute variation than the genomic background, but the ratio of within-to between-group absolute variation is especially low. The fact that these regions are consistent among all three taxa is an indication that common processes in the different greenish warbler populations have influenced these patterns.

**Fig. 6.**
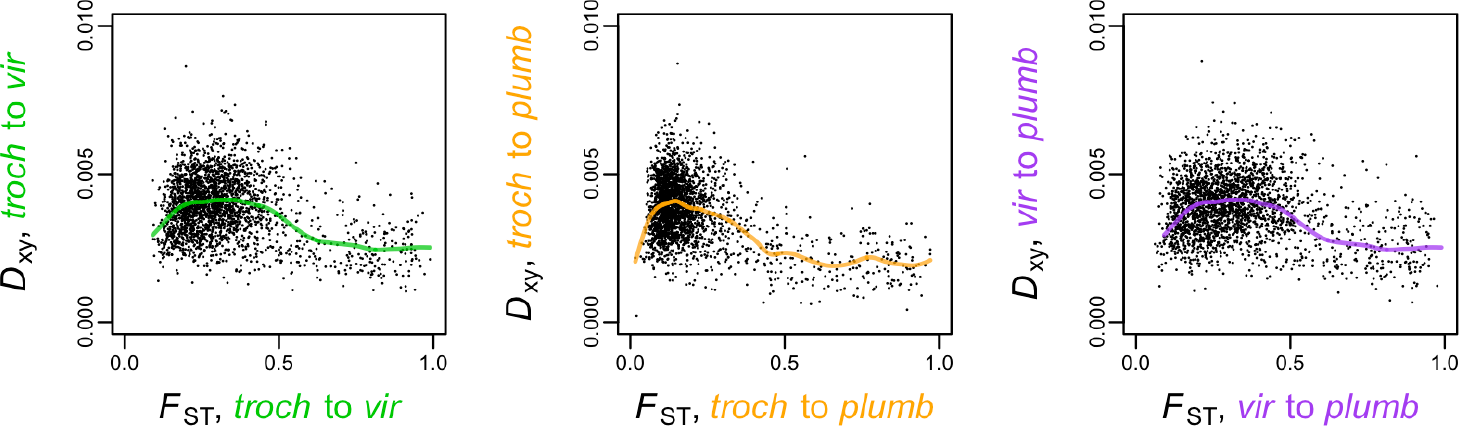
Genomic areas with high relative differentiation (*F*_ST_) tend to have low absolute nucleotide differentiation (*D*_xy_). Each graph shows *D*_xy_ versus *F*_ST_ between two greenish warbler subspecies. Each dot represents a single autosomal window containing 5000 nucleotide sites. Each comparison shows a similar pattern of a small subset of the 2486 autosomal windows deviating strongly from the majority, having large relative differentiation (right side of graph) and low absolute variation (low on the graph). Colored lines show the cubic splines fit of *D*_xy_ to *F*_ST_ (with smoothing parameter equal to one, using the “smooth.spline” function in R). The correlation among all windows is significantly negative in each pair of populations (Spearman’s rank correlation: *troch* to *vir*, *r*_s_= −0.115, P = 9.4*10^−9^; *troch* to *plumb*, *r*_s_ = −0.239, P < 10^−15^; *vir* to *plumb*, *r*_s_ = −0.052, P < 9.6*10^−3^).

**Fig. 7.**
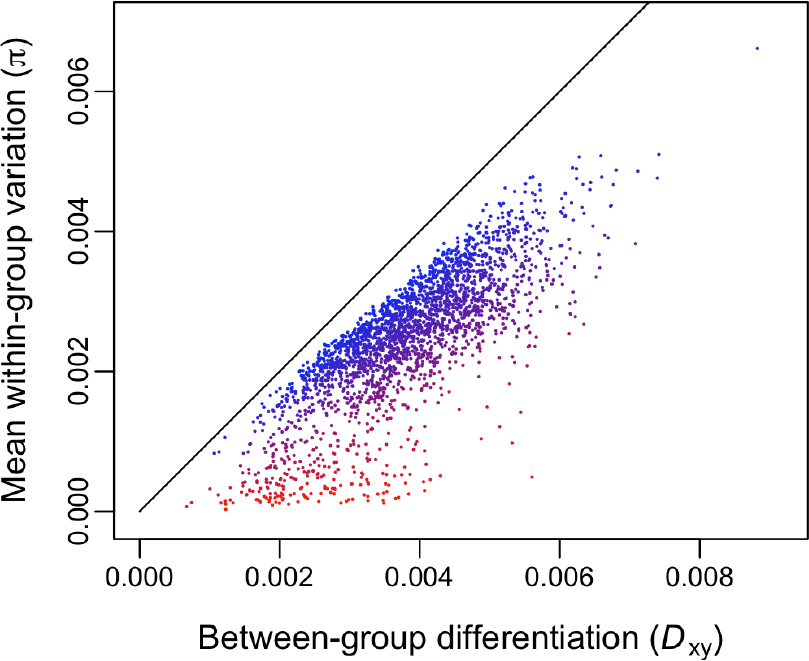
Autosomal windows with high relative differentiation (*F*_ST_; illustrated with increasing red color, whereas blue indicates low *F*_ST_) tend to have low between-group absolute differentiation (*D*_xy_) and exceptionally low average within-group variation (π). This graph shows the comparison of *viridanus* to *plumbeitarsus* (15 individuals each), but all other comparisons of greenish warbler populations show similar patterns. The diagonal line shows the 1:1 relationship that would be expected if within-group variation matched between-group differentiation (i.e., with no population differentiation).

**Fig. 8.**
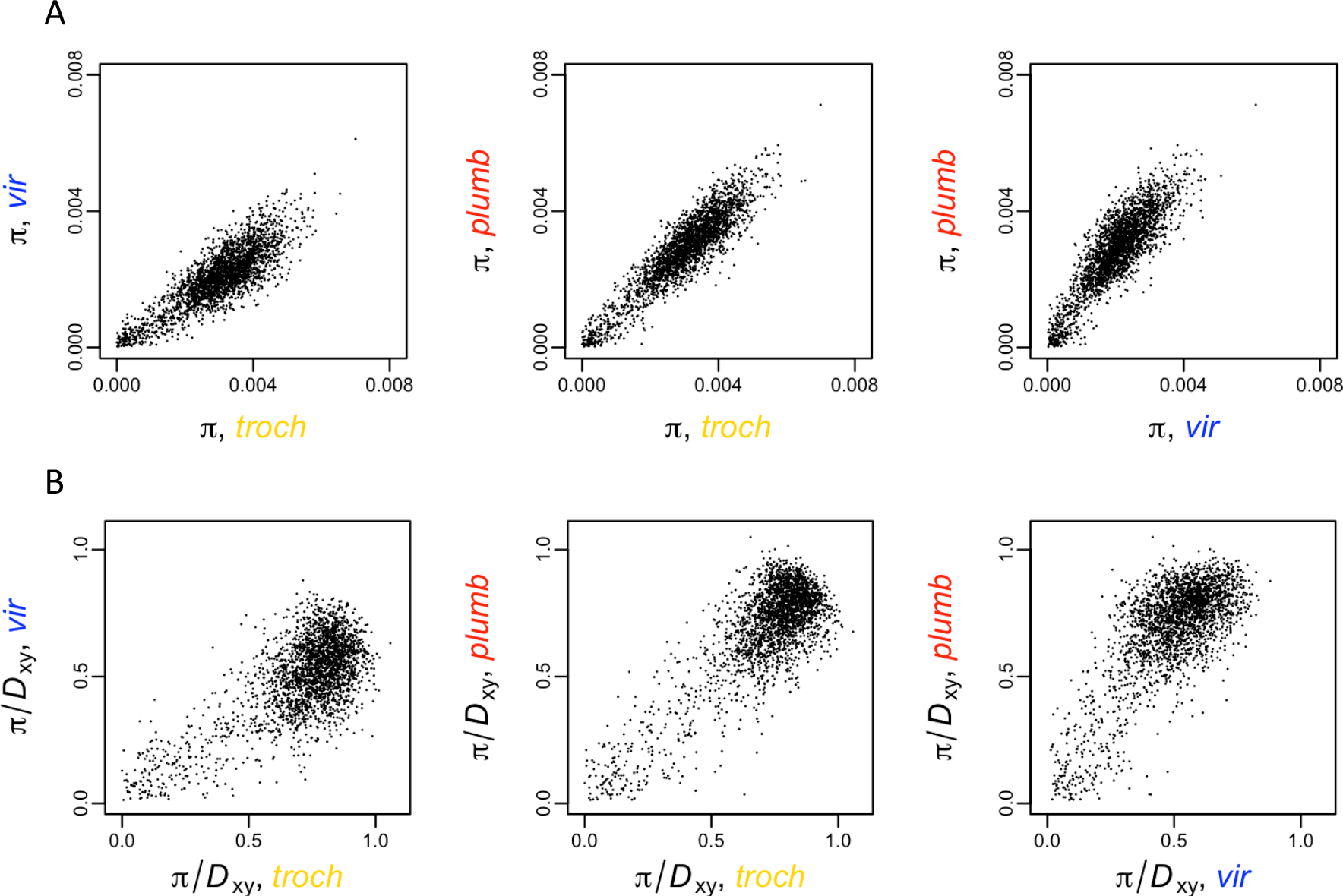
The three major greenish warbler taxa show consistent pattern of which windows show low within-group nucleotide diversity (π; top row of graphs), even when standardized by between-group absolute differentiation (i.e., the ratio of π to maximum between-group nucleotide diversity; bottom row). Each plot shows the relationship among autosomal windows of π of one taxon to that of another (top), or windowed within-group nucleotide variation (π) divided by the maximum between-group nucleotide differentiation (*D*_xy_) out of all three comparisons (bottom). Each dot represents a single autosomal window. Relationships are strong and highly significant (Pearson’s correlation test, with df = 2484: *troch* vs. *vir*. *r* = 0.840 [top] and 0.691 [bottom]; *troch* vs. *plumb*: *r* = 0.907 and 0.805; *vir* vs. *plumb*: *r* = 0.854 and 0.732; for each, P < 10^−15^). This analysis is based on 15 individuals per group.

We considered how variation in mutation rates across the genome might affect these patterns. Regions with reduced mutation rates are expected to show lower between-group absolute differentiation and within-group variation, assuming all else is equal. The effect should be similar (i.e., proportional) on both between-and within-group variation, hence variation in mutation rate among genomic regions would not explain the association between high relative differentiation and low absolute differentiation.

Nevertheless, to explore whether differences in mutation rate across the genome might be partly responsible for the shared patterns of variation within and between greenish warbler populations, we compared patterns in greenish warblers with those of four other related species in the same family of Phylloscopidae. To ensure no effects of differing sample sizes on our conclusions, we conducted an analysis using just a single individual from each of nine taxa (four more distantly related species of Phylloscopidae warblers, plus five greenish warbler subspecies, of which results are presented here for three; see Methods for details and justification of why a single sample per taxon is sufficient for this analysis). This analysis identified 11,055,883 invariant and 448,392 variant nucleotide sites among the nine taxa.

We reasoned that, if areas of consistently low absolute differentiation between pairs of greenish warblers are largely explained by consistently low mutation rates in those regions over evolutionary time, then we would see a strong correlation across the genome between absolute differentiation between greenish warbler populations and absolute nucleotide differentiation between distantly related species. We see evidence for only a weak correlation. For example, absolute nucleotide differentiation between *Phylloscopus fuscatus* and *Seicercus whistleri*, two of the most distantly related species in our study, shows only weak correlation with absolute differentiation between greenish warbler subspecies (e.g., compared to *trochiloides-viridanus*: *r* = 0.14, P = 4.7*10^−6^), explaining less than 2% of the variation (Fig. S2). In contrast, absolute differentiation between different pairs of greenish warbler populations is dramatically higher (e.g., *trochiloides-viridanus* compared to *trochiloides-plumbeitarsus: r* = 0.79, P < 10^−15^), explaining 62% of the variation (Fig. S2). We conclude that the strikingly similar patterns of variation across the greenish warbler genome in different populations cannot be explained as a result of phylogenetically stable differences in mutation rate across the genome.

Given the strong genomic patterns observed within the greenish warbler complex (genomic regions of consistently low within-group variation, moderately low between-population absolute nucleotide differentiation, and high relative differentiation), we asked whether another species complex within the same genus displayed similar patterns. *Phylloscopus inornatus* and *humei* are sister species that were only recently recognized to be distinct (Irwin et al. 2001a). Like the greenish warblers, these two sister species show a strong correlation among genomic regions in within-taxon diversity (Fig. 9). However, there is only a weak correlation between the two species complexes in their within-taxon diversity among genomic windows (Fig. 9), and when this within-taxon diversity is standardized by the between-taxon differentiation within each species complex (controlling for the fact that each taxon necessarily tends to have less diversity at each window than the entire species complex has at that window), the correlation is even weaker (Fig. S3), explaining only 2.2% of the variation. These results suggest that the two species complexes differ strongly in the genomic positions where factors such as low recombination rate and strong selection have caused unusually low within-group variation. Despite this difference, both complexes show similar patterns of regions of high *F*_ST_ having moderate or low *D*_xy_ and exceedingly low π (Fig. S4).

**Fig. 9.**
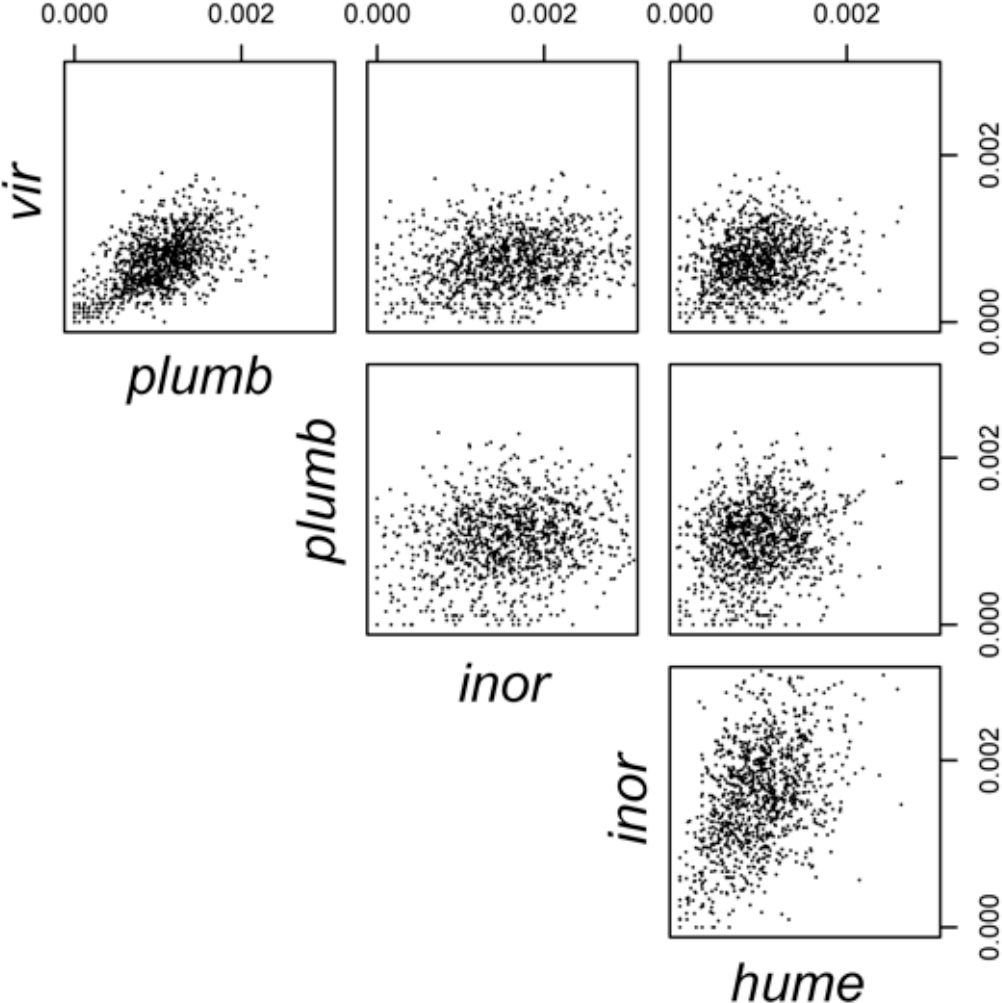
Within-group nucleotide variation (π) per window is highly correlated between taxa within the greenish warbler complex (and also within the *inornatus*/ *humei* complex), but only weakly correlated between these groups. Based on one individual per taxon, correlations of within-group nucleotide variation (π) among autosomal windows are moderately high within each species complex (e.g., *viridanus* vs. *plumbeitarsus*: *r* = 0.539, P < 10^−15^; *inornatus* vs. *humei*: *r* = 0.413, P < 10^−15^; in each, df = 1088) but much lower between these complexes (e.g., *viridanus* vs. *inornatus*: *r* = 0.189, P = 3.3 *10^−10^; the other three comparisons have similar correlations). Note that for inornatus, a few windows have π values that are too large to be shown on the plots.

Turning to the Z chromosome, recall from above that this chromosome shows higher average relative differentiation between greenish warbler populations than the autosomal genome does. To investigate whether this could be due in part to a higher rate of substitution on the Z chromosome, we compared absolute differentiation in the Z and the autosomes between distant species (between *fuscatus* and *whistleri*; and between *trochiloides* and *whistleri*). Results in both cases showed quite similar distributions of absolute differentiation in the two chromosome classes (Fig. S5); the first species pair showed no significant difference between the distribution of *D*_xy_ in the Z and autosomes (t-test; *t* = −1.83, df = 2283, P = 0.068), and the second rather surprisingly showed a slightly *lower* mean *D*_xy_ in the Z (0.0136) than in the autosomes (0.0148) (*t* = 3.27, df = 2283, P = 0.0011), suggesting a slightly lower substitution rate in the Z.

A graph of absolute differentiation between *viridanus* and *plumbeitarsus* vs. mean within-group variation (Fig. 10) shows that this sex chromosome shows similar patterns as other high– *F*_ST_ regions of the genome: *D*_xy_ and π are both significantly low compared to most of the autosomal genome (see stats in caption to Fig. 10). Despite this overall pattern, no windows on the Z chromosome reach the exceedingly low levels of π seen in some parts of the autosomal genome (Fig. 10).

**Fig. 10.**
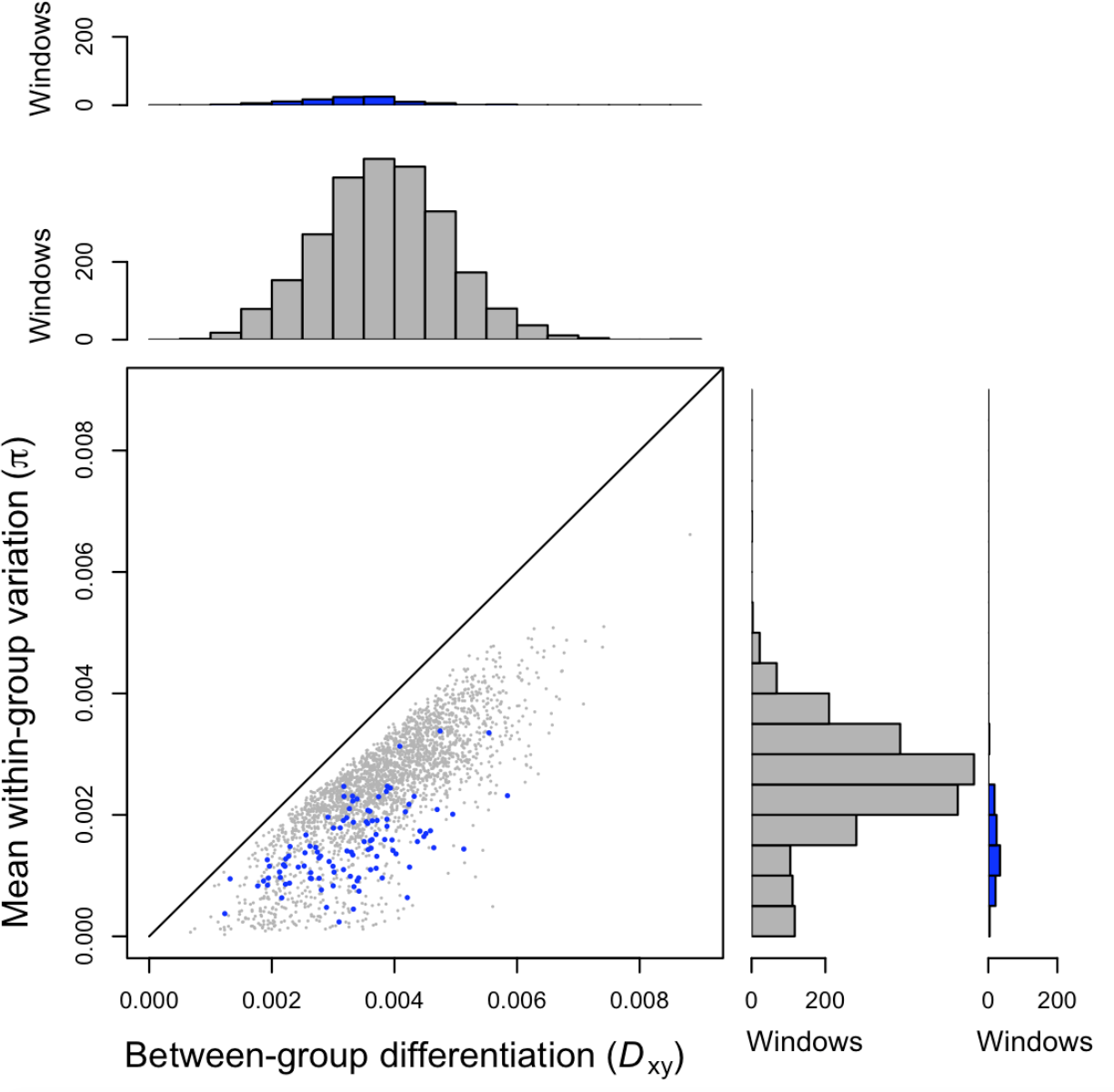
The Z chromosome (blue) differs strongly from the rest of the genome (grey) in the distribution of absolute nucleotide differentiation (*D*_xy_) and mean within-group nucleotide differentiation (π). This figure shows the comparison of *viridanus* to *plumbeitarsus* (15 individuals each), with each dot representing a single window of 5000 nucleotide sites; all other comparisons of greenish warbler populations show similar patterns. The diagonal line shows the 1:1 relationship that would be expected if within-group variation matched between-group differentiation (i.e., with no population differentiation). Histograms along each axis show that the Z-chromosome has lower *D*_xy_ and π than the rest of the genome (Welch’s t-test; *D*_xy_: *t* = 5.90, df = 115.2, P = 3.7*10^−8^; π: *t* = 15.68, df = 124.6, P < 10^−15^).

## Discussion

Prior knowledge regarding phenotypic variation among greenish warblers led us to expect some combination of parallel and non-parallel patterns of genomic change among greenish warbler populations. But given that even parallel phenotypic changes can in theory be brought about through different changes at the level of the genome, we began this study with the expectation that patterns of genomic differentiation would be highly idiosyncratic, with each population showing its own peculiar patterns in terms of which genome regions show reduced or inflated within-and between-group variation. In striking contrast to this expectation, regions with reduced within-group nucleotide diversity are remarkably similar in the three focal populations, and these regions are of high between-group relative differentiation between all pairs of populations. Moreover, these regions of high relative differentiation tend to have surprisingly low absolute differentiation between populations. Overall, these patterns indicate remarkable commonalities in the genomic regions that are subject to recurrent selection in diverse populations of greenish warblers.

By examining how these patterns relate to those within and between more distantly related species of warblers, we have been able to ask whether they may partially be due to factors that are structured across the genome in a relatively constant way over broad spans of evolutionary time. For instance, recombination rate (Renaut et al. 2013; Burri et al. 2015) and mutation rate might be expected to differ between different parts of the genome (due to structures such as centromeres and telomeres) in a relatively consistent way across a phylogeny. Likewise, the intensity of selection (background or directional) might vary consistently across the genome over broad spans of evolutionary time. If such phylogenetically-conserved factors have a large influence on patterns of variation in π and *F*_ST_, we would expect similar patterns of variation in these statistics across the genomes of greenish warblers and outgroup species. In contrast to this expectation, we find only weak (e.g. 1-5% variation explained) or no correlation between the structure of variation in greenish warblers and those within or between outgroup species. We conclude from this that the component of variation in these factors that is relatively constant over evolutionary time explains little of the genomic structuring of differentiation in greenish warblers. However, variation in these factors that is more localized in the phylogeny (i.e., confined to just the greenish warbler complex, due to rapid change in these factors over evolutionary time) could play an important role.

Although the location of regions of especially low within-group variation differ between the greenish warbler complex and the *P. inornatus*/ *humei*complex, the relationships between within-group variation, between-group absolute differentiation, and between-group relative differentiation are remarkably similar. In both cases, regions of high relative differentiation tend to have moderate or low absolute differentiation and very low within-group variation. These similarities point to common causal processes, although the genomic locations at which those processes are focused differ in the two species complexes.

To build an understanding of what processes may lead to the observed patterns, consider that we can view nucleotide differentiation between a pair of individuals as an estimate of the relative age of their common ancestor (Zeng and Corcoran 2015), also known as their relative coalescence time (to truly estimate time, we would need to know the mutation rate, but assuming the rate is constant allows us to estimate relative coalescence times; Fig. 2). Thus, the regions of high relative divergence have very short within-group coalescence times, and moderately short between-group coalescence times. The ratio of these between-to within-group coalescence times is high, consistent with the high relative differentiation (*F*_ST_). The rest of the genome (areas of low relative differentiation) have long coalescence times, both within-and between-population, and the ratio between them is much closer to one, implying that over much of the genome within-group common ancestors tend to be almost as old as between-group common ancestors.

Hence, the genomic regions of high relative divergence tend to be those where all greenish warblers share a common ancestor unusually recently, and where all individuals within a specific subspecies of greenish warbler are even more closely related. Nachman and Payseur (2012) and Cruickshank and Hahn (2014) proposed an explanation for such regions of low absolute differentiation and high relative differentiation between two taxa: selection (background and/or directional) in the common ancestral taxon reducing variation in those regions of the genome, and subsequent selection in both daughter taxa reducing the within-group variation even proportionally more (Fig. 2C). Delmore et al. (2015) proposed a related explanation: that certain regions experience selective sweeps that pass between geographic races within a geographically variable species complex, reducing variation in those regions dramatically compared to the rest of the genome, and recurrent selection in those regions then reduces within-group variation even more (Fig. 2D). Both models can be reasonably applied to greenish warblers.

Evaluation of whether regions of low absolute differentiation result from reduced variation in the common ancestor or more recent selective sweeps depends somewhat on evaluating the relative importance of background selection (due to deleterious mutations; Charlesworth et al. 1993) and positive selection (due to new beneficial mutations) in contributing to these patterns. Both selective forces can cause reductions in diversity in a panmictic common ancestor (Cruickshank and Hahn 2014), but only positive selection leads to selective sweeps over broad geographically-structured species. Zeng and Charlesworth (2011) presented a detailed analysis of a variety of background selection scenarios, using both structured coalescent models and forward-in-time simulations; they obtained values of *B*(*T*_2_), a measure of the diversity-reducing effect of background selection, ranging from 0.37 to 0.82 depending on the parameters used, meaning that the largest effect estimated was that background selection resulted in 37% of the diversity expected under pure neutrality, such that variability in nucleotide diversity (π) between different parts of the genome would vary at most roughly 2.7-fold (i.e., 1 / 0.37). In contrast, within each of the greenish warbler taxa, nucleotide diversity varies between different genomic windows to a much greater degree (Figs. 3, 4, 7, 8): for example, in the comparison of *trochiloides*to *viridanus*, mean n is 5.2-fold smaller (19%) in high-*F*_ST_(*F*_ST_ > 0.6) windows than in other windows (*F*_ST_ < 0.6), and 14.8-fold smaller (6.7%) in very-high-*F*_ST_(*F*_ST_ > 0.9) windows than in low-*F*_ST_windows (*F*_ST_ < 0.6). Although more modelling is needed to fully explore all the possible impacts of background selection (Zeng and Charlesworth 2011; see also Zeng and Corcoran 2015, which shows modest effect of background selection on π, *D*_xy_, and *F*_ST_), we conclude that modelling to date does not support background selection as being responsible for the very large heterogeneity in nucleotide variation across the genome of greenish warblers, although it almost certainly plays some role in these patterns.

Given the above, we conclude that beneficial mutations followed by selective sweeps are likely involved in generating patterns of variation in greenish warbler genomes. Such sweeps appear to have occurred within each subspecies (explaining regions of low π) as well as more deeply in time (explaining regions of low *D*_xy_), either in a panmictic common ancestor or across a geographically variable ancestor. Given the evidence for long-standing geographic structure in greenish warblers (e.g. deep mitochondrial phylogeny, phenotypic and genomic differentiation, large geographic range over a geographically complex continent) and the evidence for some current gene flow throughout the whole species complex (Irwin et al. 2001b, 2005; Alcaide et al. 2014), we find a model in which selective sweeps moved throughout a geographically structured ancestor to be most parsimonious. This could perhaps be investigated further by using the genomic variation along with inferred mutation rates to infer the age of a putative panmictic common ancestor, and then examining whether the distribution of coalescent times for windows across the genome (as inferred from *D*_xy_) is consistent with coalescence of all windows in that common ancestor (i.e., the model in Fig. 2C has a tighter range of coalescence times than the model in Fig. 2D). Such an analysis would need to make a variety of assumptions and is beyond the scope of this paper, but we reason that such a model of coalescence in a panmictic common ancestor is difficult to reconcile with the broad distribution of observed *D*_xy_ values (Fig. 7), as the common ancestor would need to be simultaneously very recent and very large. This reasoning leads us to invoke selective sweeps through a geographically structured ancestor as the most parsimonious history for regions with the lowest *D*_xy_ values.

We propose that the results are most consistent with a model in which recurrent selection, gene flow, and partial reproductive isolation likely play important roles. In this combined model, gene flow among geographically differentiated populations allows global selective sweeps to occur at specific regions where globally favorable mutations have arisen (for an example of such a sweep of a genomic region between two mosquito species, see Norris et al. 2015; and see Staubach et al. 2012 for evidence of sweeps between subspecies of House Mouse). When a genomic region undergoes a geographically global selective sweep, both between-population and within-population variation in that region is greatly reduced compared to the genomic background. Subsequent mutation and selection within each population then reduce the within-group variation at those regions even more, and increase relative differentiation between populations. If the fixation of these subsequent mutations is due in part to local adaptation that differs between populations, these regions might then play a role in (partial) reproductive isolation by causing reduced fitness in hybrids. If so, regions that play a role in reproductive isolation (i.e., contain “speciation genes”) do not tend to have high absolute differentiation—this is because they in fact have a more recent common ancestor than the rest of the genome.

We call this model the “sweep-before-differentiation model” (Fig. 2D) for the formation of peaks of high relative differentiation. It is particularly applicable to situations in which there is a low and/or intermittent gene flow between populations, in contrast with other models in which gene flow is either high (Fig. 2A) or zero (Fig. 2B). The low gene flow prevents homogenization of neutral or locally-adapted parts of the genome, but facilitates the spread of globally advantageous mutations. The genomic regions that undergo global sweeps can then diverge through selection within each local population, possibly leading to those regions causing low fitness in hybrids. This model is completely compatible with the idea that the effects of selection are strongest when it occurs on (multiple linked) genes in areas of low recombination (Noor and Bennett 2009; Nachman and Payseur 2012; Renaut et al. 2013; Burri et al. 2015).

This “sweep-before-differentiation model” for the formation of peaks of high relative differentiation incorporates the somewhat counterintuitive idea that all individuals within a species complex are more closely related in the differentiation peaks than they are elsewhere in the genome (Delmore et al. 2015). Because of the historical emphasis in the literature on relative differentiation (*F*_ST_) being directly related to gene flow between populations (reviewed by Whitlock and McCauley 1999), such peaks have often been interpreted as regions that are more distantly related between populations compared to elsewhere in the genome. However, most of the theory relating *F*_ST_ to gene flow is based on assumptions of selective neutrality. When selection plays an important role, the relationship between *F*_ST_ and gene flow can be much more complex (Whitlock and McCauley 1999). Patterns of absolute differentiation reveal that, in fact, regions of high relative differentiation are likely areas that share a more recent common ancestor, suggesting there may have been more recent gene flow in those regions than in the rest of the genome, which may share high levels of variation due to shared ancestral polymorphism rather than recent gene flow.

A large literature has discussed the commonly observed pattern of greater *F*_ST_ in the Z chromosome than on autosomes (Charlesworth et al. 1987; Ellegren et al. 2012; reviewed by Oyler-McCance et al. 2015). We note that our results are not consistent with one of the commonly proposed explanations—that there is a higher mutation rate (and therefore substitution rate) on the Z chromosome, due to the higher proportion of time that Z chromosomes occur in males (where mutation rates have been proposed to be higher). Rather, our results indicate a similar distribution of *D*_xy_ for the Z chromosome and the autosomal genome between distant species of warbler in our analysis, and between greenish warblers a lower distribution of *D*_xy_ was observed in the Z chromosome than in autosomes. The latter observation, along with the very low within-group variation at the Z chromosome (and concomitant high relative differentiation) is consistent with the well-known hypothesis (Charlesworth et al. 1987; Ellegren et al. 2012; Oyler-McCance et al. 2015) that the Z chromosome is particularly prone to recurrent selective episodes. In this respect, the entire Z chromosome displays similar characteristics of differentiation as the “islands of relative differentiation” in the autosomal genome, suggesting the Z chromosome may also be subject to geographic sweeps followed by local reductions of variation. The lower effective population size of the Z compared to autosomes (Charlesworth et al. 1987) and an expected lower recombination rate because it is hemizygous in females (Qvarnström and Bailey 2009) could also contribute to these patterns.

We note that these inferences could not have been made without decomposing relative nucleotide differentiation (*F*_ST_) into components of between-group absolute nucleotide differentiation (*D*_xy_) and within-group nucleotide differentiation (π). We have followed the lead of Nachman and Payseur (2012) and Cruickshank and Hahn (2014) here, and we reiterate their call for close examination of these statistics in analyses of population differentiation and its causes. Furthermore, we emphasize that great insight can be gained from plotting the relationships between these variables across the genome. In particular, we suggest that researchers regularly plot the relationship (among genomic windows) between absolute differentiation and average within-group variation (Fig. 7). In the case of no selection and panmixia (that is, high migration between populations), points should be clustered near the 1:1 line (that is, variation within each group is similar to variation between). Population differentiation will move the genome away from (to the right of) this line, as between-group variation becomes greater than within-group variation. If differentiation is due only to mutation and lack of gene flow (i.e., without selection), the points will move gradually away from the 1:1 line. If selection reduces within-population variation in some parts of the genome, these regions will move steadily lower on the graph, and the position of those regions along the horizontal axis shows whether those regions have low absolute differentiation (indicating ancestral reductions in diversity of those windows) or high absolute differentiation (suggesting those parts harbor genes causing reproductive isolation, whereas the rest of the genome flows between the forms at a higher rate). The plot can also be used to examine the properties of regions of high relative differentiation, by coloring points according to *F*_ST_ (Fig. 7).

In conclusion, we see remarkable similarities in the genomic patterns of variation within and between three phenotypically divergent populations within the greenish warbler ring species. Comparisons with outgroup species suggest these similarities are not well explained by phylogenetically-conserved differences in mutation rate or recombination rate across the genome (at least in terms of the components of variation in those factors that stay relatively consistent over moderately long evolutionary time). In particular, the differences observed between the greenish warbler complex and the *Phylloscopus inornatus*/ *humei* complex in genomic locations of relative divergence cast doubt on the hypothesis that regions of high relative differentiation in these warblers correspond to genomic structural features such as centromeres or telomeres (e.g., Ellegren et al. 2012), unless those features have largely changed location in the roughly 12 million years of evolution since these two species complexes shared a common ancestor (Johansson et al. 2007; Price 2010). We conclude that the patterns are supportive of linked selection impacting most strongly on the same portions of the genome within each population of greenish warblers, but that these locations differ among species groups separated by more evolutionary time. We note however that the evidence for selection acting on similar regions does not indicate that selection pressures are identical in the three populations; rather, it simply indicates that selection is operating on traits encoded by similar regions. Even selection on the same gene could be in differing directions in two populations (e.g. selection for larger wing bars in one place, and smaller in another) while having a similar effect in causing reduced within-group variation. Given the strong evidence for certain phenotypes typically being under selection in warbler populations (e.g. beak size and shape, wing bars, migratory route and distance, singing behavior; Richman and Price 1992; Irwin 2000; Irwin and Irwin 2005; Price 2008; Tietze et al. 2015), it is reasonable that certain genomic regions also are consistently under more intense selection than others. The evidence that these regions have lower between-group coalescence times suggests that there have been selective sweeps in some of these regions throughout the species complex (whether in the common ancestor or more recently). The possibility that regions of high relative differentiation have experienced more recent gene flow than other genomic regions has received very little previous consideration, and could dramatically alter our understanding of the process of genomic differentiation during speciation. Altogether, the results of this study point to a remarkable level of repeatability of patterns of selection and genomic differentiation within a widespread and geographically variable species complex.

## Acknowledgements

This work was supported by the Natural Sciences and Engineering Research Council of Canada (Discovery Grant number 311931) and a Marie Curie International Outgoing Fellowship within the 7th European Community Framework Programme (project no. 273773). Assistance in the field was provided by Z. Benowitz-Fredericks, J. Gibson, S. Gross, G. Kelberg, A. Knorre, K. Marchetti and B. Sheldon. For additional samples we thank P. Alström, K. Marchetti, U. Olsson, T. Price, A. Richman, E. Scordato, and J. Tiainen. The study benefitted from discussions and/or comments on the manuscript from R. Burri, B. Harr, J. Mallet, T. Price, L. Rieseberg, D. Schluter, D. Toews, S. Wang, C. Wei, and M. Whitlock.

## Data Accessibility

– Whole-genome shotgun sequencing reads archived at NCBI SRA: *viridanus*: SRX472921; *trochiloides*: SRX1625194; *plumbeitarsus*: SRX1630030.
– GBS reads archived at NCBI SRA: SRX473141.
– We are currently working on submitting the three assembled individual genomes to NCBI Genbank. We will archive the consensus genome and vcf files containing called genotypes in the Dryad Digital Repository following acceptance of the manuscript.

## Author Contributions

DEI, MA, and JHI designed the research; MA conducted the laboratory work; KED assembled the genomes; KED and GLO assisted with the bioinformatics pipeline; DEI designed the R scripts, conducted the analysis of GBS reads, and wrote the paper with input from all authors.

## Supporting Information

To accompany this manuscript: Irwin DE, Alcaide M, Delmore KE, Irwin JH, and Owens GL. Recurrent selection explains parallel evolution of genomic regions of high relative but low absolute differentiation in greenish warblers. Submitted to *Molecular Ecology*.

**Table S1.** Summary statistics for de novo genome assemblies obtained with SOAPdenovo, including the number of ultra-conserved elements (UCEs) from Faircloth et al. 2012 that aligned to each assembly. UCEs were identified using whole genome alignments of chicken, anole and zebra finch. The broad set of UCEs included 5561 elements; the narrow set was limited to UCEs with higher coverage and included 2560 elements.

| Taxon | Length (bp) | # sequences | # Ns in sequence | N50 | GC % | # UCEs (broad) | # UCEs (narrow) |
| --- | --- | --- | --- | --- | --- | --- | --- |
| *P. t. viridanus* | 1,011,681,890 | 223,054 | 7,147,412 | 20,406 | 41.46 | 5101/5561 | 2551/2560 |
| *P. t. trochiloides* | 1,011,245,339 | 413,267 | 9,021,417 | 7,319 | 41.58 | 5081/5561 | 2548/2560 |
| *P. t. plumbeitarsus* | 997,759,019 | 670,969 | 8,367,615 | 3,515 | 41.72 | 5104/5561 | 2554/2560 |

**Table S2.** Locations of regions that were locally re-aligned with Mauve (Darling et al. 2010).

| Chromosome | Starting position (bp) | End position (bp) | Chromosome length (bp) | Seed weight used |
| --- | --- | --- | --- | --- |
| 1A | 43,312,085 | 43,494,011 | 76,983,199 | 5 |
| 2 | 36,314,908 | 36,536,933 | 106,153,545 | 5 |
| 2 | 102,036,725 | 102,595,309 | 106,153,545 | 5 |
| 3 | 44,894,889 | 45,059,641 | 115,734,465 | “auto” |
| 3 | 83,248,205 | 84,192,906 | 115,734,465 | “auto” |
| 6 | 36,207,248 | 36,264,701 | 37,369,755 | 5 |
| 10 | 14,057,576 | 14,113,875 | 21,311,711 | “auto” |
| 12 | 3,497,365 | 5,226,257 | 21,841,389 | 5 |

**Table S3.**
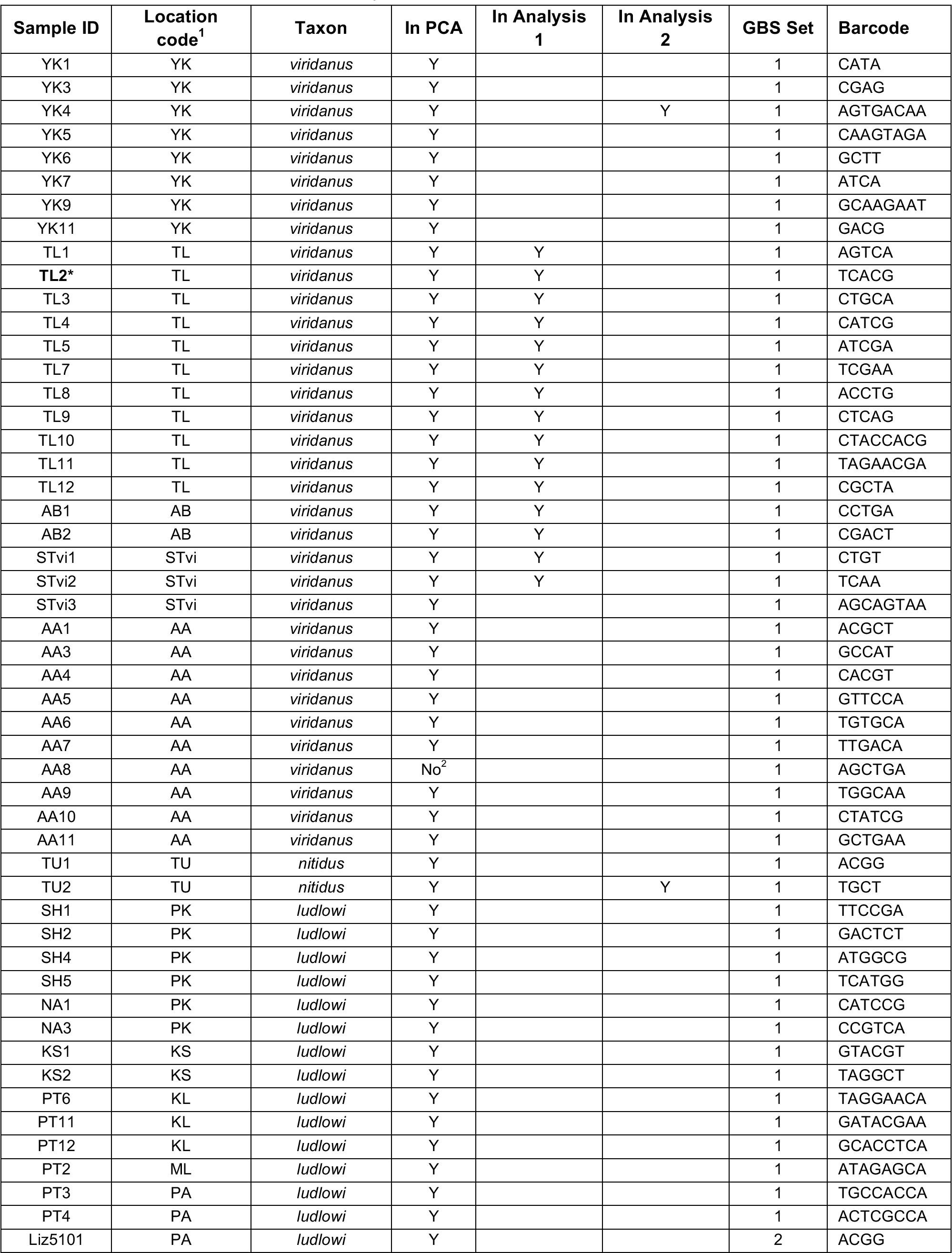
Individuals used in the study.

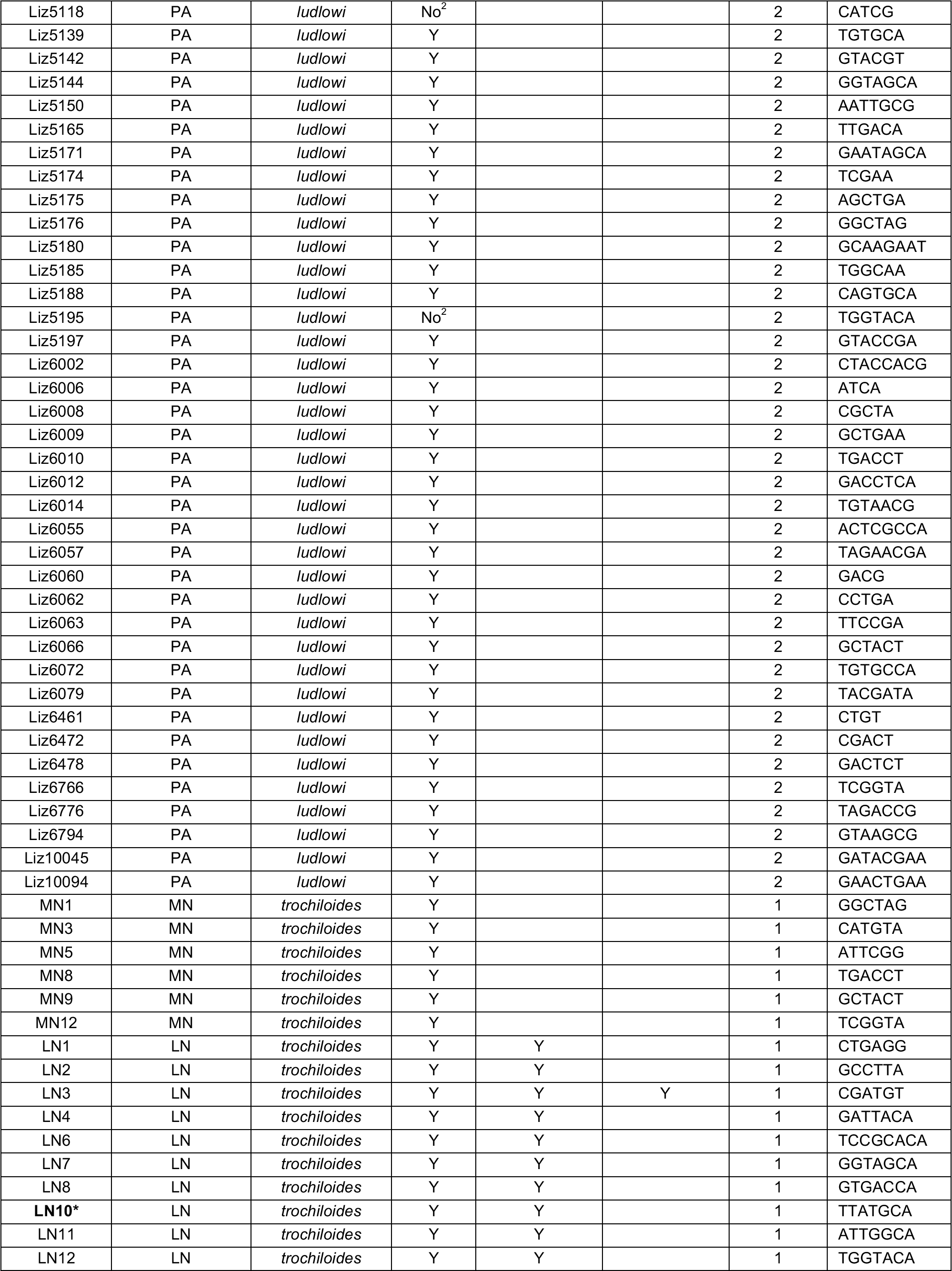

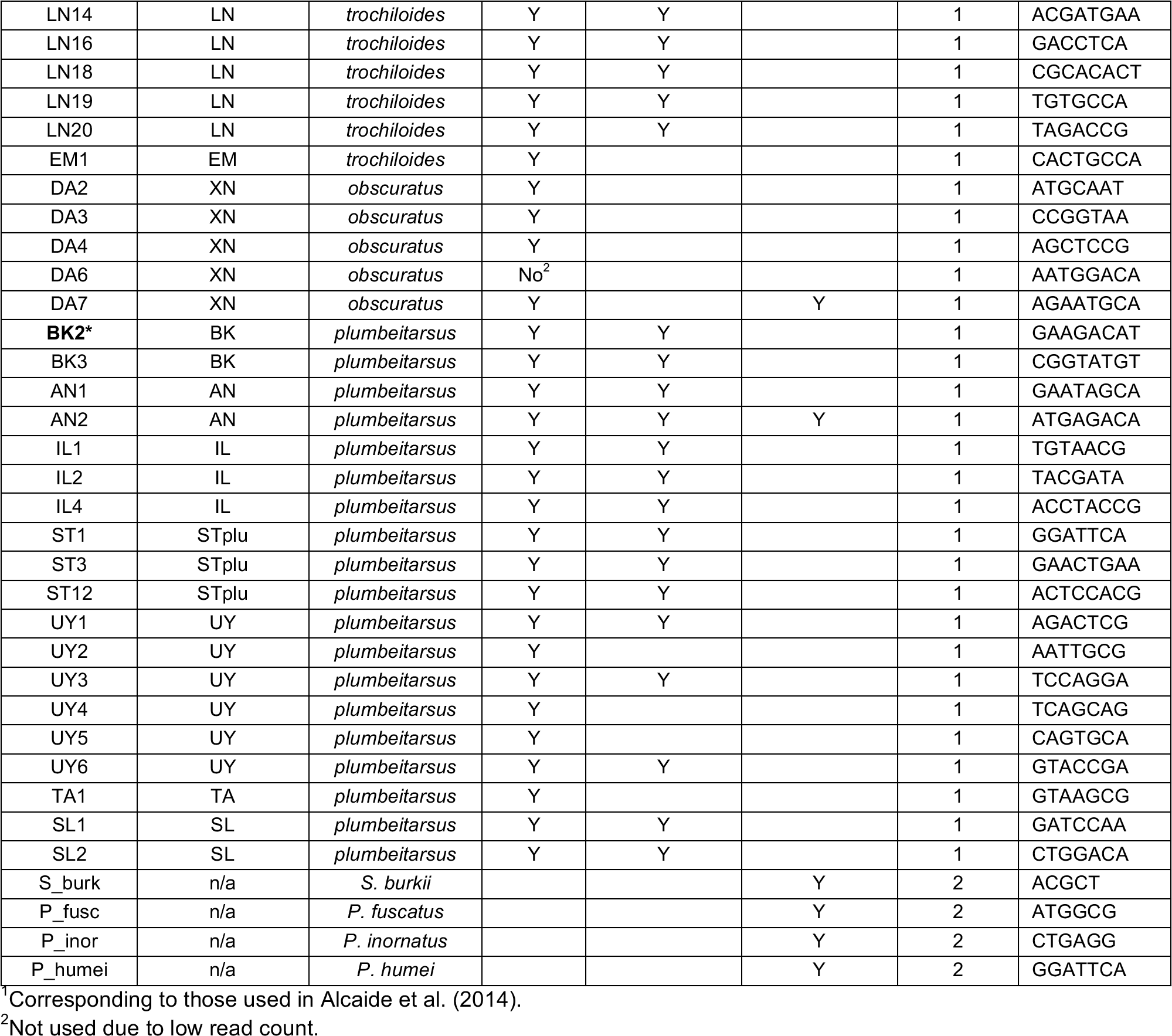

**Fig. S1.**
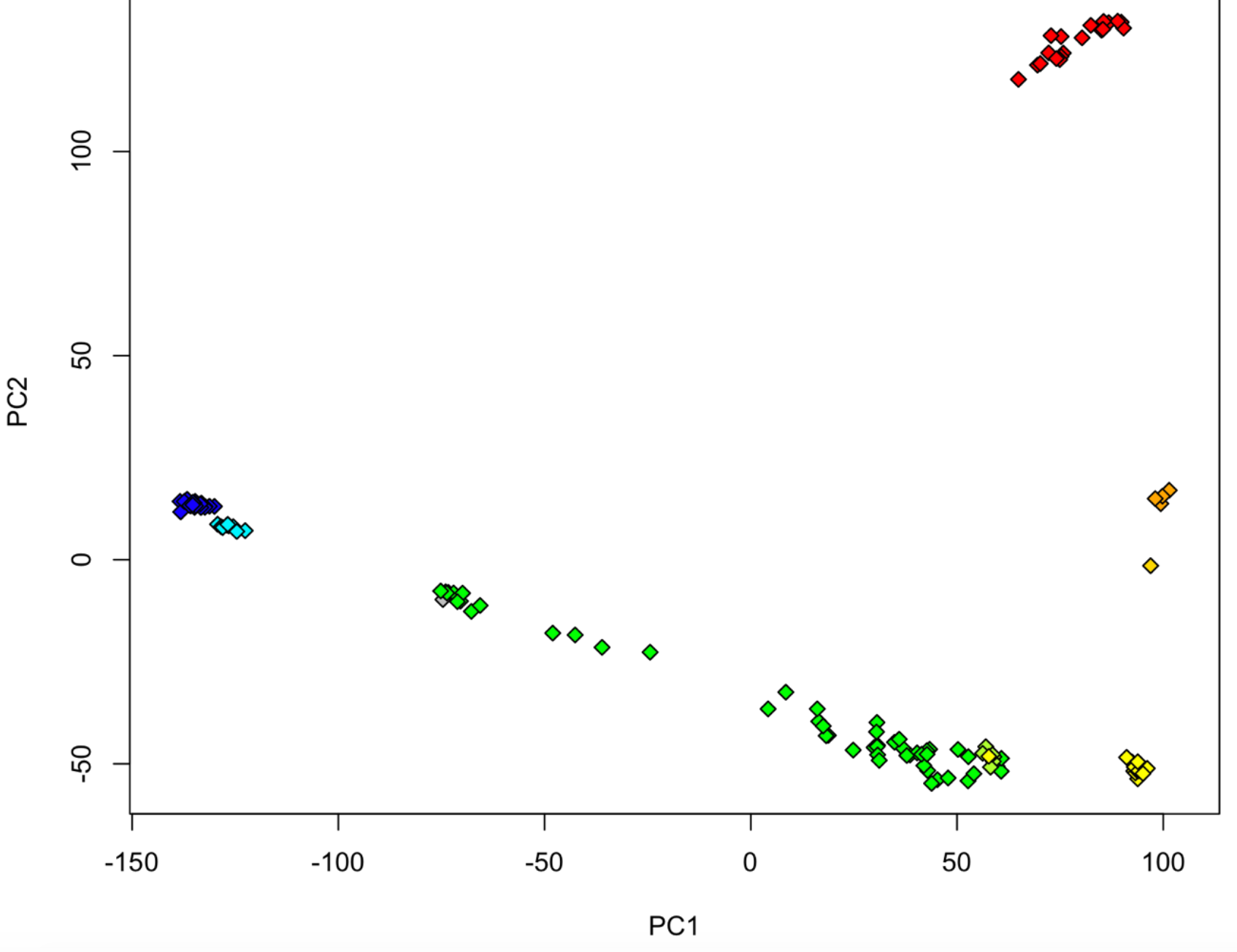
Genomic relationships around the ring of greenish warblers. Results of a principal components analysis based on 580,356 SNPs among 135 individual greenish warblers. Each symbol represents a single individual, with colors indicating geographic region (i.e., subspecies identity; See Figure 1 and Table S3). Points close together have similar genomic signatures, whereas those far apart differ strongly. The first principal component explains 15.0% of genomic variation, and the second explains 6.6%.

**Fig. S2.**
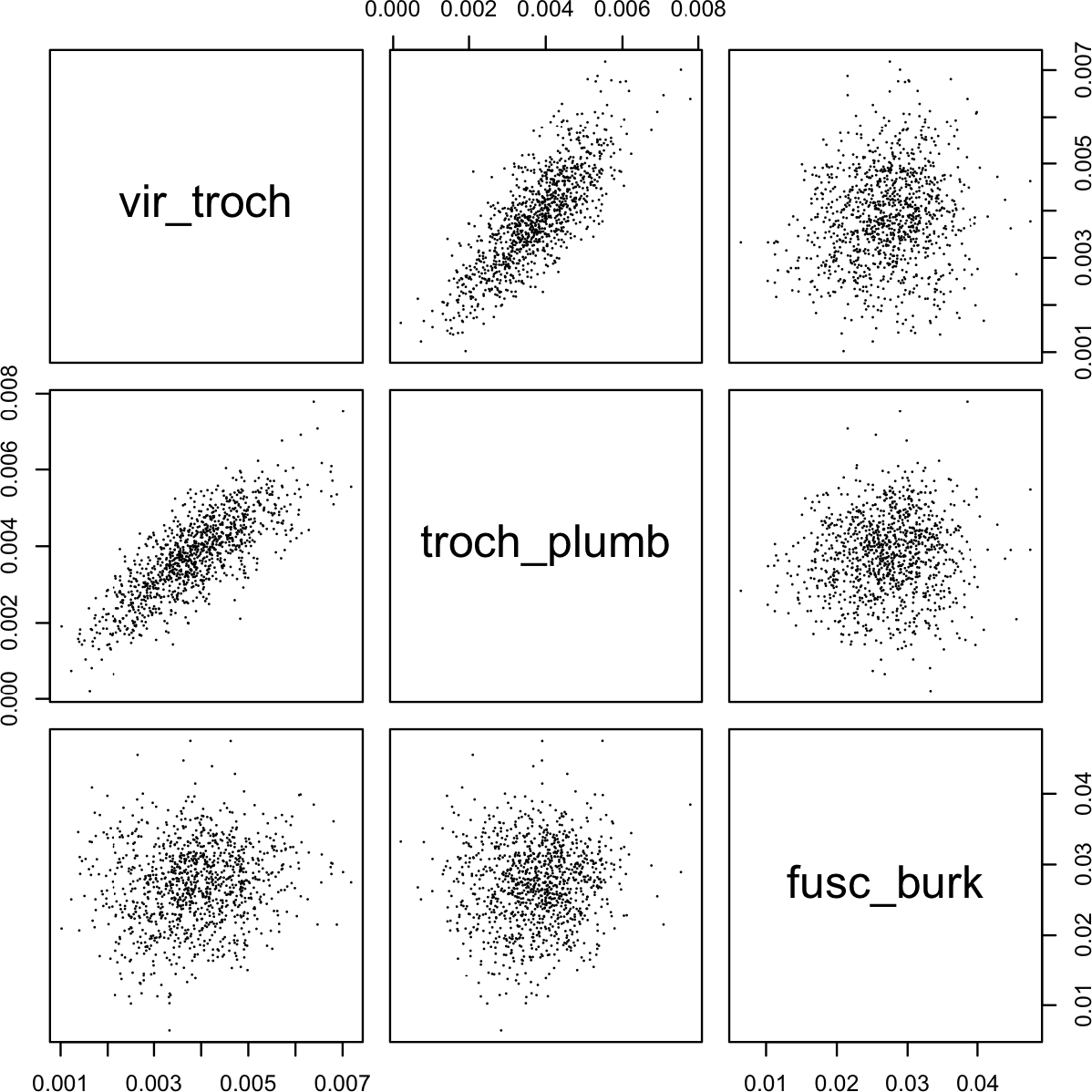
Levels of absolute differentiation (*D*_xy_) within genomic windows are highly correlated between comparisons of pairs of greenish warbler populations (e.g., *trochiloides-viridanus* vs. *trochiloides-plumbeitarsus*, shown here), but only weakly correlated between each pair of greenish warbler populations and the distantly related species *Phylloscopus fuscatus* and *Seicercus burkii*. Each of the correlations is statistically significant, but only the comparisons within greenish warblers explain a large portion of variation (e.g., 62% in the *trochiloides-viridanus* vs. *trochiloides-plumbeitarsus* comparison; Pearson’s correlation, with df = 1088: *r* = 0.787, P < 10^−15^). Genetic distance among windows between *fuscatus* and *burkii* explains less than 2% of the variation among windows in absolute nucleotide differentiation between pairs of greenish warblers (*trochiloides*-*viridanus* vs. *fuscatus-burkii*: *r*= 0.138, P = 4.7*10^−6^; *trochiloides-plumbeitarsus* vs. *fuscatus-burkii*: *r* = 0.079, P = 0.0091). These results are based on the analysis of variation in a single individual of each taxon (see Methods).

**Fig. S3.**
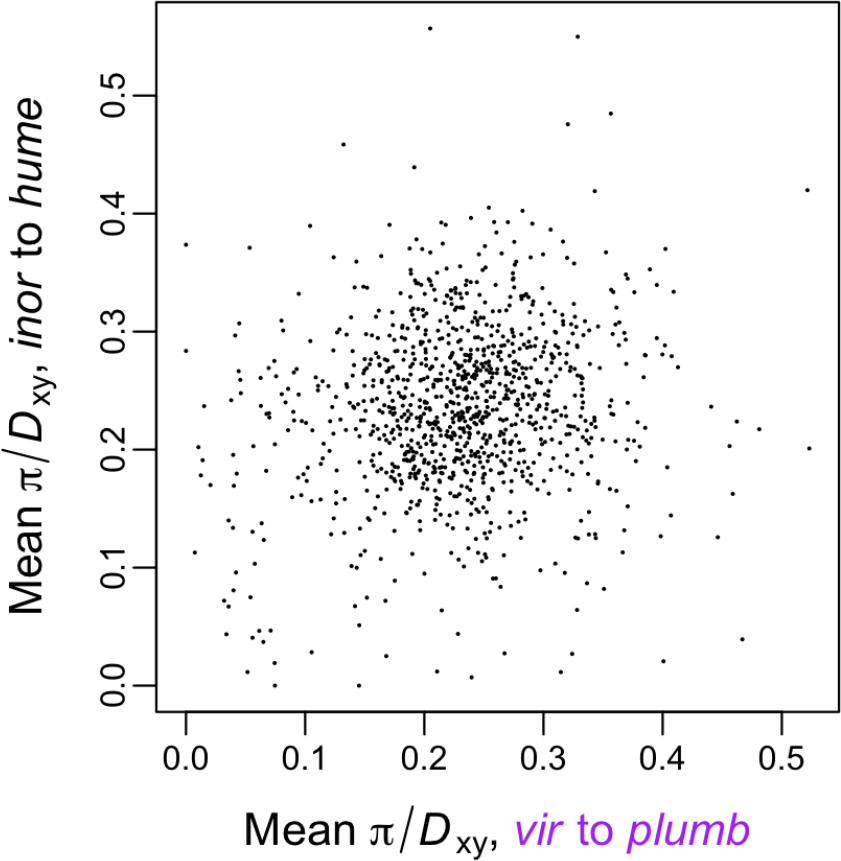
The ratio of within- to between-group nucleotide variation per window is only weakly correlated between the greenish warbler complex and the *inornatus / humei* complex. Each axis represents the mean standardized within-group variation of two taxa (the average within-group nucleotide variation in two taxa divided by the absolute nucleotide differentiation between them) within a single species complex. Greenish warblers (*specifically*, *viridanus* and *plumbeitarsus*) are on the horizontal axis, and *inornatus*and *humei* are on the vertical. There is only a weak correlation, explaining 2.2% of the variation (*r* = 0.149, df = 1088, P = 8.0*10^−7^).

**Fig. S4.**
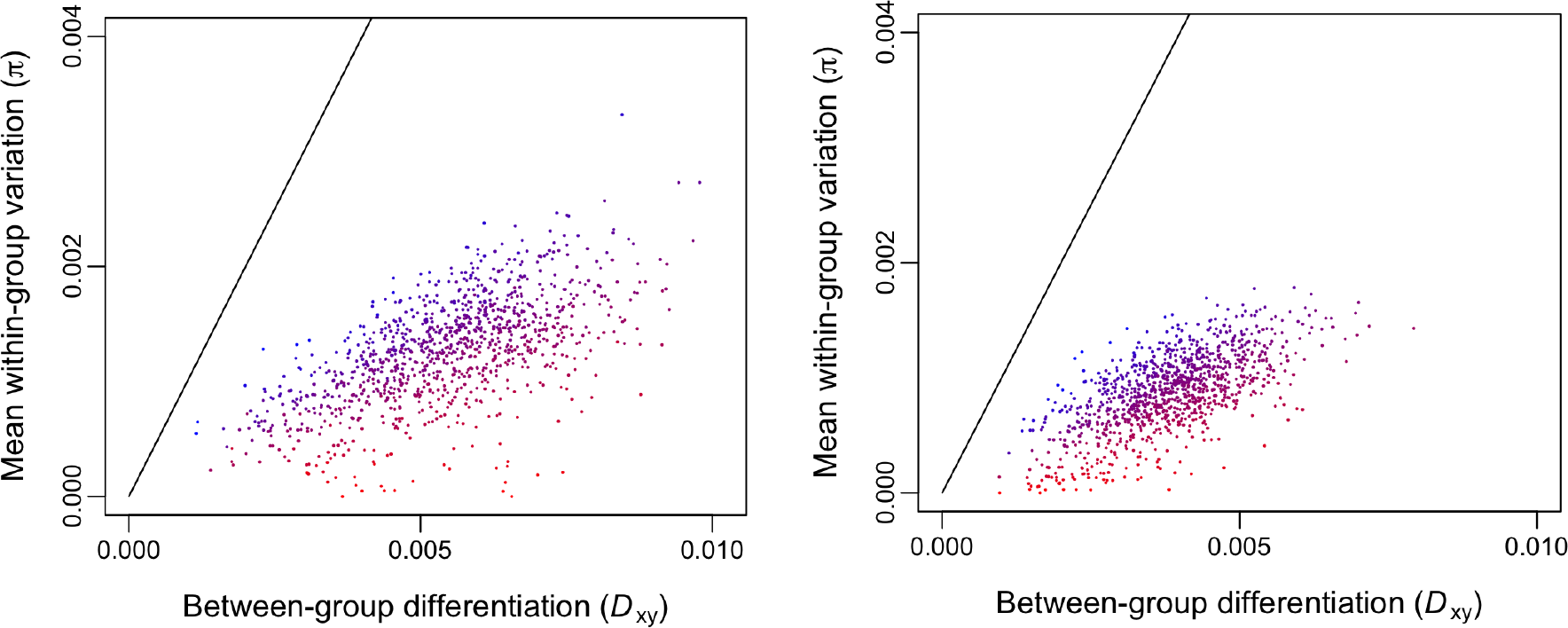
Analyses of two phylogenetically distant taxon pairs of *Phylloscopus*warblers (left: *humei / inornatus*; right: *viridanus / plumbeitarsus*) shows that in both, autosomal windows with high relative differentiation (*F*_ST_; illustrated with increasing red color, whereas blue indicates low *F*_ST_) tend to have moderate or low between-group absolute differentiation (*D*_xy_) and exceptionally low average within-group variation (π). These graphs are based on a single individual per taxon. Note that within-group variation is greatly underestimated in this analysis (in contrast to the 15-individual-per-taxon analysis presented in Fig. 7), due to the small sample size. This effect is similar for all windows, such that comparisons among windows within an analysis are valid.

**Fig. S5.**
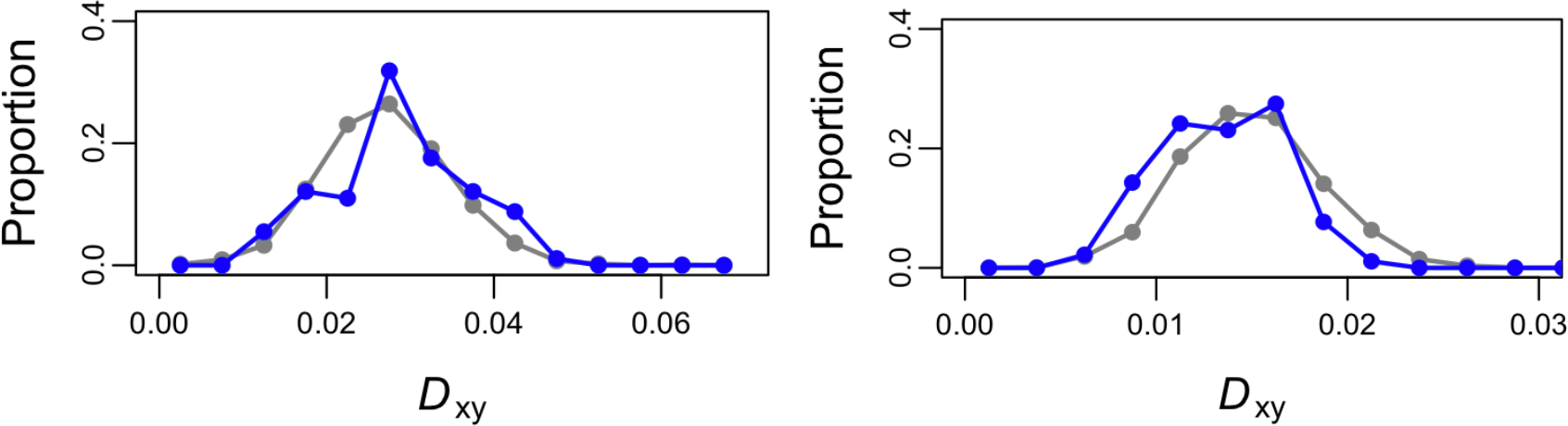
The distribution of absolute nucleotide differentiation (*D*_xy_) is similar in the Z chromosome (blue) and the autosomal genome (grey) in comparisons between distantly related species (left: *fuscatus* and *burkii*; right: *trochiloides* and *burkii*). There is no significant difference between the distribution of *D*_xy_in the Z and autosomes in the comparison between *fuscatus* and *burkii*(t-test; *t* = −1.83, df = 2283, P = 0.068), and in the *trochiloides* - *burkii* comparison there is a slightly lower mean *D*_xy_ in the Z (0.0136) than in the autosomes (0.0148) (*t*= 3.27, df = 2283, P = 0.0011).

## References

AlcaideM, ScordatoESC, PriceTD, IrwinDE (2014). Genomic divergence in a ring species complex. Nature, 511, 83–85.

BauerB, GokhaleCS (2015) Repeatability of evolution on epistatic landscapes. Scientific Reports, 5, 9607.

BolgerAM, LohseM, UsadelB (2014) Trimmomatic: a flexible trimmer for Illumina sequence data. Bioinformatics, 30, 2114–2120.

BradburdGS, RalphPL, CoopGM (2016) A spatial framework for understanding population structure and admixture. PLoS Genetics, 12, e1005703.

BurriR, NaterA, KawakamiT, Mugal, CF, OlasonPL, SmedsL, SuhA, DutoitL, BurešS, GaramszegiLZ, HognerS, MorenoJ, QvarnströmA, Ružic, SxtreS-A, SxtreG-P, TorokTorok, EllegrenH (2015) Linked selection and recombination rate variation drive the evolution of the genomic landscape of differentiation across the speciation continuum of Ficedula flycatchers. Genome Research, 25, 1656–1665.

CainAJ (1954) Animal Species and their Evolution. Hutchinson House London.

CarneiroM, AlbertFW, AfonsoS, PereiraRJ, BurbanoH, CamposR, Melo-FerreiraJ, Blanco-AguiarJA, VillafuerteR, NachmanMW, GoodJM, FerrandN (2014) The genomic architecture of population divergence between subspecies of the European rabbit. PLoS Genetics, 10, e1003519.

CharlesworthB, CoyneJA, BartonNH (1987) The relative rates of evolution of sex chromosomes and autosomes. American Naturalist, 130, 113–146.

CharlesworthCharlesworth, MorganMorgan, CharlesworthD (1993) The effect of deleterious mutations on neutral molecular variation. Genetics, 134, 1289―1303.

ColosimoPF, HosemannKE, BalabhadraS, VillarrealJr.G, DicksonM, GrimwoodJ, SchmutzJ, MyersRM, SchluterD, KingsleyDM (2005) Widespread parallel evolution in sticklebacks by repeated fixation of ectodysplasin alleles. Science, 307, 1928–1933.

ConteGL, ArnegardME, PeichelCL, SchluterD2012The probability of genetic parallelism and convergence in natural populations. Proceedings of the Royal Society of London Series B, Biological Sciences, 279, 5039–5047.

CruickshankTE, HahnMW (2014) Reanalysis suggests that genomic islands of speciation are due to reduced diversity, not reduced gene flow. Molecular Ecology, 23, 3133–3157.

DanecekP, AutonA, AbecasisG, AlbersCA, BanksE, DePristoMA, HandsakerR, LunterG, MarthG, SherryST, McVeanG, DurbinR, 1000 Genomes Project Analysis Group(2011) The variant call format and VCFtools. Bioinformatics, 27, 2156–2158.

DarlingAE, MauB, PernaNT (2010) progressiveMauve: multiple genome alignment with gene gain, loss and rearrangement. PLoS ONE, 5, e11147.

DelmoreKE, HübnerS, KaneNC, SchusterR, AndrewRL, CamaraCamara, GuigoGuigo, IrwinDE (2015) Genomic analysis of a migratory divide reveals candidate genes for migration and implicates selective sweeps in generating islands of differentiation. Molecular Ecology, 24, 1873–1888.

EllegrenH (2013) The evolutionary genomics of birds. Annual Review of Ecology, Evolution, and Systematics, 44, 239–259.

EllegrenH, SmedsL, BurriR, OlasonPI, BackströmA, KawakamiT, KünstnerA, MäkinenH, Nadachowska-BrzyskaK, QvarnströmA, UebbingS, WolfJBW (2012) The genomic landscape of species divergence in Ficedula flycatchers. Nature, 491, 756–760.

ElshireH, GlaubitzL, SunQ, PolandJA, KawamotoK, BucklerES, MitchellSE (2011) A robust, simple genotyping-by-sequencing (GBS) approach for high diversity species. PLoS ONE, 6, e19379.

FairclothBC, McCormackJE, CrawfordNGet al.(2012) Ultraconserved elements anchor thousands of genetic markers spanning multiple evolutionary timescales. Systematic Biology, 61, 717–726.

FederJL, NosilP (2010) The efficacy of divergence hitchhiking in generating genomic islands during ecological speciation. Evolution, 64, 1729–1747.

GouldS (1990) Wonderful Life: The Burgess Shale and the Nature of History. W. W. Norton and Company, New York.

HarrB (2006) Genomic islands of differentiation between house mouse subspecies. Genome Research, 16, 730–737.

IrwinDE (2000) Song variation in an avian ring species. Evolution, 54, 998–1010.

IrwinDE (2012) Culture in songbirds and its contribution to the evolution of new species. In: Creating Consilience: Integrating the Sciences and the Humanities (eds. after edsSlingerlandE, CollardM), pp. 163–178. Oxford University Press, Oxford

IrwinDE, AlströmP, OlssonU, Benowitz-FredericksZM (2001a) Cryptic species in the genus Phylloscopus. Ibis, 143, 233–247.

IrwinDE, BenschJH, PriceTD (2001b) Speciation in a ring. Nature, 409, 333–337.

IrwinDE, BenschS, IrwinJH, PriceTD (2005) Speciation by distance in a ring species. Science, 307, 414–416.

IrwinDE, IrwinJH (2005) Siberian migratory divides: the role of seasonal migration in speciation. In: Birds of Two Worlds: the Ecology and Evolution of Migration(eds. GreenbergR, MarraPP), pp. 27–40. Johns Hopkins University Press, Baltimore, Maryland.

IrwinDE, IrwinJH, PriceTD (2001c) Ring species as bridges between microevolution and speciation. Genetica, 112–113, 223–243.

IrwinDE, ThimganMP, IrwinJH (2008) Call divergence is correlated with geographic and genetic distance in greenish warblers (Phylloscopus trochiloides): a strong role for stochasticity in signal evolution? Journal of Evolutionary Biology, 21, 435–448.

JohanssonUS, AlströmP, OlssonU, EricsonPGP, SundbergP, PriceTD (2007) Build-up of the Himalayan avifauna through immigration: a biogeographical analysis of the Phylloscopus and Seicercus warblers. Evolution, 61, 324–333.

KawakamiT, SmedsL, BackströmN, HusbyA, QvarnströmA, MugalCF, OlasonP, EllegrenH (2014) A high-density linkage map enables a second-generation collared flycatcher genome assembly and reveals the patterns of avian recombination rate variation and chromosomal evolution. Molecular Ecology, 23, 4035–4058.

LiH, DurbinR (2009) Fast and accurate short read alignment with Burrows-Wheeler transform. Bioinformatics, 25, 1754–1760.

LiH, HandsakerB, WysokerA, FennellT, RuanJ, HomerN, MarthG, AbecasisG, DurbinR, 1000 Genome Project Data Processing Subgroup(2009) The Sequence Alignment/Map (SAM) format and SAMtools. Bioinformatics, 25, 2078–2079.

LobkovskyAE, KooninEV (2012) Replaying the tape of life: quantification of the predictability of evolution. Frontiers in Genetics, 3, 246.

LuoR, LiuB, XieY, LiZ, HuangW, YuanJ, et al.(2012) SOAPdenovo2: an empirically improved memory-efficient short-read de novo assembler. Gigascience, 1, 18.

MayrE (1942) Systematics and the Origin of Species. Dover Publications, New York.

McKennaA, HannaM, BanksE, SivachenkoA, CibulskisK, KernytskyA, GarimellaK, AltshulerD, GabrielS, DalyM, DePristoMA (2010) The Genome Analysis Toolkit: A MapReduce framework for analyzing next-generation DNA sequencing data. Genome Research, 20, 1297–1303.

MeyerJR, AgrawalAA, QuickRT, DobiasDT, SchneiderD, LenskiRE (2010) Parallel changes in host resistance to viral infection during 45,000 generations of relaxed selection. Evolution, 64, 3024–3034.

MeyerJR, DobiasDT, WeitzJS, BarrickJE, QuickRT, LenskiRE (2012) Repeatability and contingency in the evolution of a key innovation in phage lambda. Science, 335, 428–432.

NachmanMW, PayseurBA (2012) Recombination rate variation and speciation: theoretical predictions and empirical results from rabbits and mice. Philosophical Transactions of the Royal Society of London. Series B, Biological Sciences, 367, 409–421.

NadeauNJ, WhibleyA, JonesRT, DaveyJW, DasmahapatraKK, BaxterSW, QuailMA, JoronM, ffrench-ConstantRH, BlaxterML, MalletJ, JigginsCD (2012) Genomic islands of divergence in hybridizing Heliconius butterflies identified by large-scale targeted sequencing. Philosophical Transactions of the Royal Society of London. Series B, Biological Sciences, 367, 343–353.

NoorMAF, BennettSM (2009) Islands of speciation or mirages in the desert? Examining the role of restricted recombination in maintaining species. Heredity, 103, 439–444.

NorrisLC, MainBJ, LeeY, CollierTC, FofanaA, CornelAJ, LanzaroGC2015Adaptive introgression in an African malaria mosquito coincident with the increased usage of insecticide-treated bed nets. Proceedings of the National Academy of Sciences of USA, 112, 815–820.

NosilP, FederJL (2012) Genomic divergence: causes and consequences. Philosophical Transactions of the Royal Society of London B, 367, 332–342.

NosilP, FunkDJ, Ortiz-BarrientosD (2009) Divergent selection and heterogeneous genomic divergence. Molecular Ecology, 18, 375–402.

Oyler-McCanceSJ, CornmanRS, JonesKL, FikeJA (2015) Z chromosome divergence, polymorphism and relative effective population size in a genus of lekking birds. Heredity, 115, 452–459.

PriceTD (2008) Speciation in Birds. Roberts and Co.Boulder, Colorado.

PriceTD (2010) The roles of time and ecology in the continental radiation of the Old World leaf warblers (Phylloscopus and Seicercus). Philosophical Transactions of the Royal Society of London B, 365, 1749–1762.

QvarnströmA, BaileyRI (2009) Speciation through evolution of sex-linked genes. Heredity, 102, 4–15.

R Core Team(2014) R: A Language and Environment for Statistical Computing. R Foundation for Statistical Computing, Vienna, Austria. URL http://www.R-project.org/

RenautS, GrassaCJ, YeamanS, MoyersBT, LaiZ, KaneNC, BowersJE, BurkeJM, RiesebergLH (2013) Genomic islands of divergence are not affected by geography of speciation in sunflowers. Nature Communications, 4, 1827.

RenautS, OwensGL, RiesebergLH (2014) Shared selective pressure and local genomic landscape lead to repeatable patterns of genomic divergence in sunflowers. Molecular Ecology, 23, 311–324.

RichmanAD, PriceTD (1992) Evolution of ecological differences in the Old World leaf warblers. Nature355, 817–821.

StaubachF, LorencA, MesserPW, TangK, PetrovDA, TautzD (2012) Genome patterns of selection and introgression of haplotypes in natural populations of the House Mouse (Mus musculus). PLoS Genetics, 8, e1002891.

TicehurstCB 1938 A Systematic Review of the Genus Phylloscopus. Trustees of the British Museum, London.

TietzeDT, MartensJ, FischerBS, SunY-H, Klussmann-KolbA, PackertM (2015) Evolution of leaf warbler songs (Aves: Phylloscopidae). Ecology and Evolution, 5, 781–798.

TravisanoM, MongoldJ A, BennettAF, LenskiRE (1995) Experimental tests of the roles of adaptation, chance, and history in evolution. Science, 267, 87–90.

TurnerTL, HahnMW (2010) Genomic islands of speciation or genomic islands and speciation?Molecular Ecology, 19, 848–850.

TurnerTL, HahnMW, NuzhdinSV (2005) Genomic islands of speciation in Anopheles gambiae. PLoS Biology, 3, e285.

ViaS (2012) Divergence hitchhiking and the spread of genomic isolation during ecological speciation-with-gene-flow. Philosophical Transactions of the Royal Society of London. Series B, Biological Sciences, 367, 451–460.

WarrenWC, ClaytonDF, EllegrenH, ArnoldAP, HillierLW, KünstnerA, et al. (2010) The genome of a songbird. Nature, 464, 757–762.

WeirBS, CockerhamCC (1984) Estimating F-statistics for the analysis of population structure. Evolution, 38, 1358–1370.

WhitlockMC, McCauleyDE (1999) Indirect measures of gene flow and migration: FST ≠1/4Nm+1). Heredity, 82, 117–125.

WichmanHA, BadgettMR, ScottLA, BoulianneCM, BullJJ (1999) Different trajectories of parallel evolution during viral adaptation. Science, 285, 422–424.

WoodTE, BurkeJM, RiesebergLH (2005) Parallel genotypic adaptation: when evolution repeats itself. Genetica, 123, 157–170.

WuC (2001) The genic view of the process of speciation. Journal of Evolutionary Biology, 14, 851–865.

XuH, LuoX, QianJ, PangX, SongJ, QianG, ChenJ, ChenS (2012) FastUniq: a fast de novo duplicates removal tool for paired short reads. PLoS ONE, 7, e52249.

ZhangG, LiB, LiC, GilbertM, JarvisED, WangJ, The Avian Genome Consortium(2014) Comparative genomic data of the avian phylogenomics project. GigaScience, 3, 26.

ZengK, CharlesworthB (2011) The joint effects of background selection and genetic recombination on local gene genealogies. Genetics, 189, 251–266.

ZengK, CorcoranP (2015) The effects of background and interference selection on patterns of genetic variation in subdivided populations. Genetics, 201, 1539–1554.

